# The RNA binding protein DAZL functions as repressor and activator of maternal mRNA translation during oocyte maturation

**DOI:** 10.1101/598805

**Authors:** Cai-Rong Yang, Gabriel Rajkovic, Enrico Maria Daldello, Xuan G. Luong, Jing Chen, Marco Conti

## Abstract

Deleted in azoospermia like (DAZL) is an RNA-binding protein playing critical function during gamete development. In fully-grown oocytes, DAZL protein is detected in prophase and levels increase four to five fold during reentry into the meiotic cell cycle. Here, we have investigated the functional significance of this DAZL accumulation in maturing oocytes. Oocyte depletion of DAZL prevents progression to MII. This maturation block is associated with widespread disruption in the pattern of maternal transcripts loading on ribosomes and their translation measured using a RiboTag IP/RNASeq or qPCR strategy. In addition to decreased ribosome loading of a subset of transcripts, we found that DAZL depletion causes also translational activation of distinct subset of mRNAs. DAZL binds to mRNAs whose translation is both repressed and activated during oocyte maturation. Unexpectedly, DAZL depletion also causes increased ribosome loading of a subset of mRNAs in quiescent GV-arrested oocytes. This dual role of repression and activation is recapitulated by using YFP reporters including the 3’UTR of DAZL targets. Injection of recombinant DAZL protein in DAZL-depleted oocytes rescues the translation of these targets as well as maturation to MII. Mutagenesis of putative DAZL-binding sites in these candidate mRNAs mimics the effect of DAZL depletion. These findings demonstrate that DAZL regulates translation of maternal mRNAs in mature oocytes, functioning both as translational repressor and activator.

## Introduction

In both males and females of most species, production of gametes is a developmental process that spans embryonic, fetal, and postnatal life and is essential for the transfer of genetic information to the progeny^1^. Germline lineage specification, expansion of the gamete precursors (PGCs), meiosis, and terminal differentiation into functional gametes are all elaborate processes that require extensive regulation of gene expression^1, 2^. Together with unique transcriptional and epigenetic mechanisms, regulation of translation plays a critical role in differentiation of these germ cells^3, 4, 5^.

In a mature mRNA, several domains play a critical role in the regulation of translational efficiency and stability^6^. These include the CAP region of the mRNA, the 5’ and 3’ UTR, and the poly(A) tail^6, 7, 8^. Translation of an mRNA is modulated through the interaction of numerous proteins with these domains of a mRNA. Indeed the assembly of ribonucleoprotein (RNPs) modulates every aspect of mRNA translation and stability. A subgroup of RNA-binding protein (RBPs) interacts with the 3’UTR of an mRNA. These RBPs participate in the formation of RNPs that are critical for the control of translational efficiency, stability and localization of the mRNAs^9^. These properties place these RBPs in a critical role in the control of protein synthesis. Among the several RBPs uniquely expressed in the germ line is the family of the Daz RNA-binding proteins^10, 11^. DAZ, DAZL and BOULE are germ-cell specific RBPs essential for gametogenesis from worms to humans^11^.

DAZL KO prevents differentiation of PGCs^10, 12, 13^. It has been proposed that DAZL functions as a translational activator by recruiting poly(A) binding proteins, which in turn promotes and stabilize interaction with the cap of mRNA, a loop conformation thought to promote stability and translational efficiency^14^. However, additional studies in PGCs suggest a repressive function for this RBP in the control of zebrafish embryogenesis and in the mouse^15, 16^. Moreover, depletion of DAZL during spermatogenesis has been associated with mRNA destabilization^17^.

We have previously reported that acute depletion of DAZL from fully grown mouse oocytes using morpholino oligonucleotides causes disruption of the progression through meiosis^18, 19^. Here we have used this *in vitro* model to define the pattern of translation dependent on the function of this RBP. We find that DAZL depletion causes both increases and decreases in translational efficiency of a wide range of transcripts expressed in the oocyte and these effects are reversible and recapitulated by regulation of reporter translation of candidate DAZL targets.

## Results

### DAZL is expressed in fully grown oocytes and is depleted upon inhibition of DAZL mRNA translation

We have reported that DAZL protein is detectable in fully grown GV-arrested oocytes with protein levels increasing further up to MII ^18^. This finding is at odd with data of others where DAZL protein was only borderline detectable or could not be detected^20, 21^. Here, we have re-evaluated the expression of DAZL during oocyte maturation using a newly developed monoclonal DAZL antibody (see ‘Materials and Methods’). Western blot analysis of extracts from oocytes at different stages of maturation (0, 2, 4, 6, 8hrs) reveals an immunoreactive polypeptide with mobility corresponding to that of DAZL (37 kDa) in GV oocytes, and a three-to-fourfold increase in protein levels up to 8 hrs of *in vitro* maturation (MI) (Fig.1a, Supplenmentary Fig. 1e), in complete agreement with our previous reports using *in vivo* matured oocytes. Note that the identity of the immunoreactive band is further confirmed by morpholino kncockdown experiments (see Fig. 1b, Supplementary Fig. 1f). Thus, two different antibodies document that DAZL is expressed in GV oocytes and that accumulation of the protein increases with maturation, conclusions consistent with our *Dazl* mRNA translation data with *in vivo* and *in vitro* matured oocytes (Supplementary Fig.1a, b, c).

**Figure 1.**
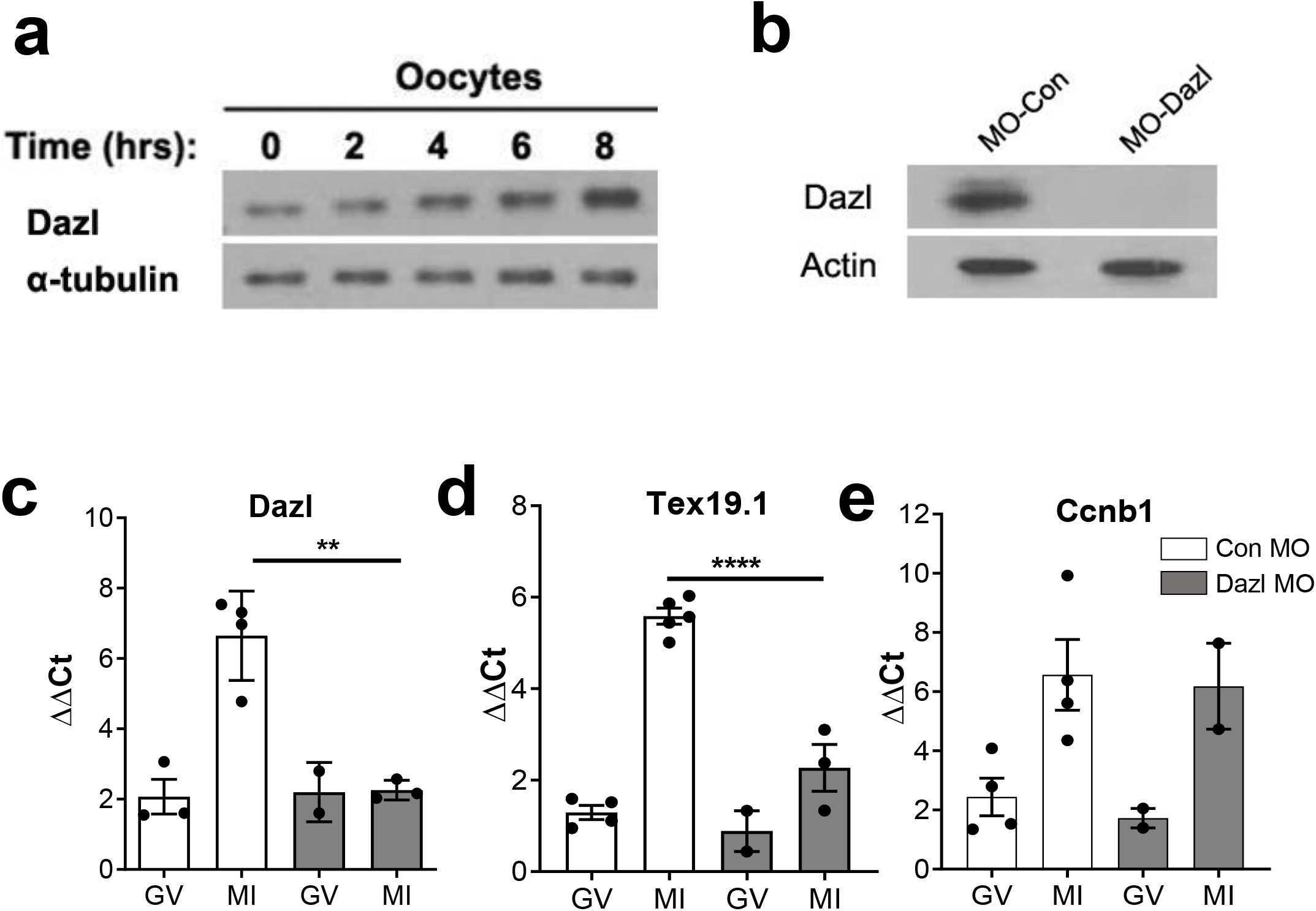
Interference with Dazl mRNA translation depletes the oocyte of the DAZL protein and inhibits translation of a specific downstream target. **(a)** DAZL protein accumulation during the transition from the GV-to-MI stage of oocyte maturation. Accumulation of α-tubulin was used as a loading control. GV stage oocytes from wild type mice were cultured *in vitro* for maturation. After 0-8 hrs maturation, oocytes were collected, lysed in sample buffer, and used for Western blot analysis. 150 oocytes per lane was loaded in this experiment. **(b)** Morpholino down-regulation of DAZL protein. GV stage oocytes from wild type or DAZL Heterozygous mice were injected with CON or DAZL MO. Oocytes were preincubated overnight with 2 μM milrinone and then cultured in inhibitor-free medium for maturation. After 6hrs, oocytes were collected and used for Western blot analysis. A representative experiment of the three performed is reported. **(c-e)** Ribosome loading of endogenous *Dazl* and *Tex19.1*, but not *CcnB1*, mRNA is blocked after DAZL depletion. GV stage oocytes from wild type or *Dazl*^+/−^ mice were injected with CON-MO or Dazl MO and preincubated overnight in 2 μM milrinone, then cultured in inhibitor-free medium for maturation. Oocytes were collected at 0 hr and 6 hrs for RiboTag IP followed by qPCR analysis. (GV:germinal vesicle; MI:Meiosis I). Each dot represents the average of triplicate measurements from independent biological samples collected in different days. ** P<0.01; **** P<0.0001.

To determine whether preventing *Dazl* mRNA translation effectively depletes the oocytes of the DAZL protein, GV-arrested oocytes from RiboTag^fl/fl^: Zp3-CRE, Dazl^+/+^ or Dazl^+/−^ mice were injected with a scrambled (Con-MO) or DAZL targeting morpholino (DAZL-MO) respectively, to maximize DAZL protein removal. Blockage of *Dazl* mRNA translation by this specific MO markedly reduces (94.235% decrease +/− 0.025, Mean +/− SEM, N=3) the endogenous DAZL protein expression compared to a CON-MO (Fig. 1b). To further assess the effectiveness and specificity of the treatment, we used RiboTag IP/qPCR to evaluate the MO effect on ribosome loading onto endogenous mRNAs during the transition from germinal vesicle stage (GV) to Meiosis I stage (MI). We observe a significant decrease in ribosome recruitment onto the *Dazl* mRNA, confirming the effectiveness of the MO in blocking initiation of translation, with consequent depletion of the protein from oocytes (Fig. 1c). TEX19.1 is an established target of DAZL^19^. We show that the *Tex19.1* mRNA loading on ribosomes is significantly decreased after injection of DAZL-MO at the MI stage (Fig. 1d). Conversely, knockdown of DAZL had no effect on ribosome loading onto the non-target *CcnB1* (Fig. 1e), which is again consistent with our previous report ^18, 19^. No detectable effect on total transcript levels was detected under these conditions. Confirming what previously reported by us, DAZL depletion disrupts oocyte maturation to MII (see below). Further control experiments where immunoprecipitation was performed with WT rather than RiboTag mice yield only background signal, confirming the specificity of the RiboTag immunoprecipitation (Supplementary Fig. 1d). These pilot experiments document that DAZL knockdown specifically disrupts DAZL target loading onto ribosomes with high selectivity since it does not affect the translation of *CcnB1*, an mRNA that does not interact with DAZL. Additionally, they confirm that RiboTag IP in oocytes depleted of DAZL is an effective strategy to assess the role of this RBP in endogenous maternal mRNA translation.

### Ribosome loading onto maternal mRNAs is disrupted in oocytes depleted of DAZL

For a genome-wide analysis of the effect of DAZL depletion on translation of oocyte endogenous mRNAs, GV oocytes from RiboTag^fl/fl^: Zp3-CRE, Dazl^+/+^ or Dazl^+/−^ mice were injected with Con-MO or DAZL-MO. After overnight recovery, oocytes were collected at 0 hr (GV) or cultured in inhibitor-free medium to mature for up to 6 hrs (MI). Although the changes in translation would be more pronounced if measured in fully matured MII oocytes, this short maturation time was selected to monitor early effects of DAZL depletion, thus, avoiding the potential confounding effects of the blockage of maturation to MII, and a potential decrease in oocyte viability. When we compare total mRNAs from CON-MO and DAZL-MO in GV-arrested oocytes (overnight incubation in PDE inhibitors), few differences are detected (Fig. 2a). Also comparison of ribosome loading in the CON-MO at 0 hr and 6 hrs shows changes in ribosome loading qualitative similar to those reported previously with polysome array or other RiboTag IP/RNASeq data sets with non injected oocytes (Supplementary Fig. 2a). However, when we compare RiboTag IP/RNASeq in the 6 hrs DAZL-MO versus 6hrs CON-MO group, we observe complex changes in maternal mRNA ribosome loading (Fig. 2b). Ribosome loading onto the majority of transcripts present in the oocyte is not significantly affected by the DAZL depletion (grey dots in Fig. 2b). However, we detect a decrease in ribosome loading onto a group of transcripts (blue dots in Fig. 2b, 551 transcripts, FDR < 0.05), a finding consistent with the theory that DAZL functions as a translational activator. Surprisingly, we also identify a group of transcripts whose translation increases (red dots in Fig. 2b, 170 transcripts, FDR < 0.05) after DAZL removal. This latter finding indicates that directly or indirectly DAZL has a role in repression of translation. As an example of the RNASeq data, ribosome loading onto *Tex19.1, Txnip, Akap10*, and *Nsf* mRNAs at 0 hr and 6 hrs of maturation are reported in Fig. 2d, 2e. These mRNAs are significantly immunoprecipitated by DAZL antibody as shown in our DAZL RIP-Chip dataset (Fig. 2c). We found no clear pattern in the changes in total mRNA levels after DAZL depletion (Fig. 2d). RiboTag IP/RNA-Seq shows an increase in ribosome loading (HA immunoprecipitation) for transcripts *Tex19.1* and *Txnip* during maturation in CON-MO injected oocytes, whereas the recruitment is blunted after DAZL KD in MI stage(Fig. 2e). Conversely, ribosome loading onto *Akap10* and *Nsf* mRNAs is increased after DAZL depletion (Fig. 2e). These data provide an initial indication that that DAZL functions not only a translational activator, but also a translational repressor during the GV-to-MI transition. Given the fact that no significant differences in total mRNA were detected between the CON-MO and DAZL-MO groups, calculation of the translational efficiency (HA-IP:input ratio) shows the same trend (Supplementary Fig. 2b).

**Figure 2.**
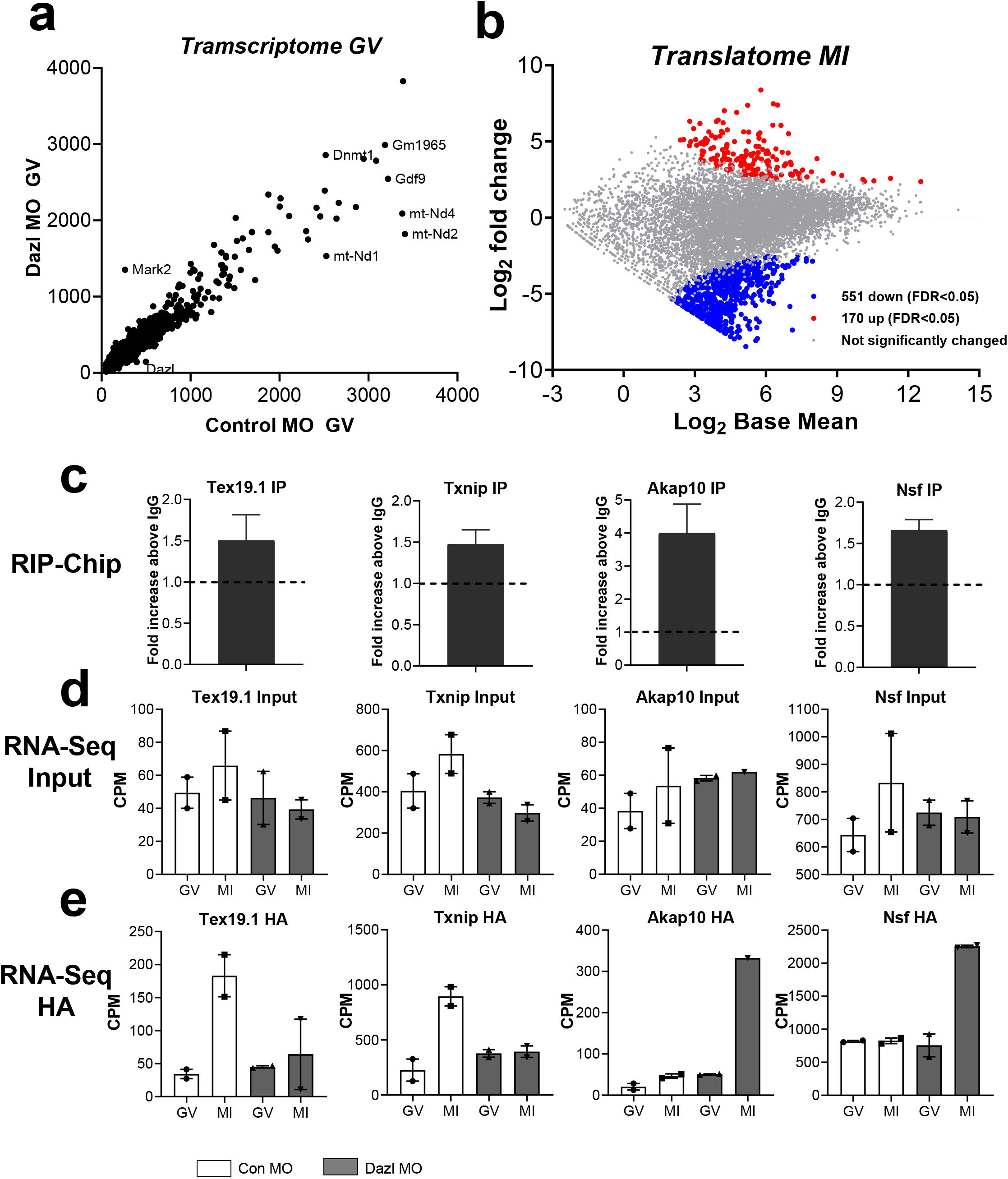
Maternal mRNA loading onto ribosome is disrupted in oocytes depleted of DAZL. **(a)** Comparison of the transcriptomes of oocytes injected with control and *Dazl* MO. Oocytes from wild type mice were injected with a CON-MO whereas oocytes from heterozygous *Dazl* mice were injected with a DAZL-MO. Oocytes were incubated overnight in the presence of milrinone and the following morning were collected for RiboTag IP/RNASeq as described in the ‘Materials and Methods’. The average input (total transcripts) CPM data from duplicate biological replicates is reporterd. **(b)** Comparison of transcripts recovered by RiboTag IP/RNASeq in Control and *Dazl* MO injected oocytes (MI). GV stage oocytes from wild type or *Dazl*^+/−^ mice were injected with CON-MO or Dazl-MO. After overnight preincubation with 2 μM milrinone, oocytes were cultured in inhibitor-free medium to allow reentry into the meiotic cell cycle. Oocytes were collected at 6hrs for RiboTag IP and RNA-Seq analysis as detailed in the methods. Ribosome loading of the majority of transcripts present in the oocyte is not significantly changed after DAZL removal (grey dots). Ribosome loading of a subgroup (551 transcripts) of mRNAs (blue, FDR < 0.05) is significantly decreased, while ribosome loading of a distinct subgroup (170) of transcripts is significantly increased after DAZL removal (red dots, FDR < 0.05). **(c-e)** effect of DAZL depletion on RNA levels and ribosome loading of representative DAZL interacting targets. **(c)** DAZL RIP-Chip of oocyte extracts immunoprecipitation of selected mRNAs is reported as the fold enrichment DAZL AB/IgG N=3. **(d and e)** GV stage oocytes from wild type or *Dazl*^+/−^ mice were injected with control or *Dazl* MOs. After overnight preincubation with 2 μM milrinone, oocytes were cultured in inhibitor-free medium for maturation. Oocytes were collected at 0hr (GV) and 6hrs (MI) for RiboTag IP and RNA-Seq analysis. **(d)** RNASeq data from supernatants (input) from RiboTag IP of control and *Dazl* MO **(e)** RiboTag IP/RNASeq analysis documented an increase in ribosome loading onto these transcripts (*Tex19.1* and *Txnip*) in control oocytes but the increase is absent after DAZL KD. *Akap10* and *Nsf* mRNA translation is increased after DAZL depletion.

### The dual effect of DAZL depletion is confirmed by RiboTag IP/qPCR

To confirm the opposing effect of DAZL depletion on translation, we performed RiboTag IP/qPCR with independent biological samples to monitor the recruitment of representative transcripts to the ribosome/translation pool. GV stage oocytes from wild type or DAZL Heterozygous mice were injected with control or DAZL MO. After overnight preincubation with 2 μM milrinone, oocytes were cultured in inhibitor-free medium. Oocytes were collected at 6 hrs for RiboTag IP /qPCR analysis. This RNA quantification confirms that the overall transcripts levels (input of RiboTag IP/qPCR) are not affected, including transcripts coding for *Dazl, Tex19.1, Txnip, Rad51C, Btg4, Ero1, Oosp1, Obox5, Ireb2*, and *Tcl1* (Supplementary Fig. 3a). However, RiboTag IP/qPCR shows a decrease translation for several candiates after DAZL removal, similar to that observed with the RiboTag IP/RNASeq (Fig. 3a, c). *Dppa3* and *CcnB1* are used as negative control, as they are not regulated by DAZL during oocyte maturation^18, 19^. Conversely, translation of transcripts coding for *Akap10, Cenpe, Nsf, Ywhaz, Nin*, and *YTHDF3* is upregulated after DAZL removal (Fig. 3b, d), while the overall transcripts levels are not changed (Supplementary Fig. 3b). *Gdf9*, used as negative control, is not affected by DAZL depletion. These results not only confirm our RiboTag IP/RNASeq data, but also indicate that DAZL may plays dual function (both translational activator and repressor) during oocyte maturation.

**Figure 3.**
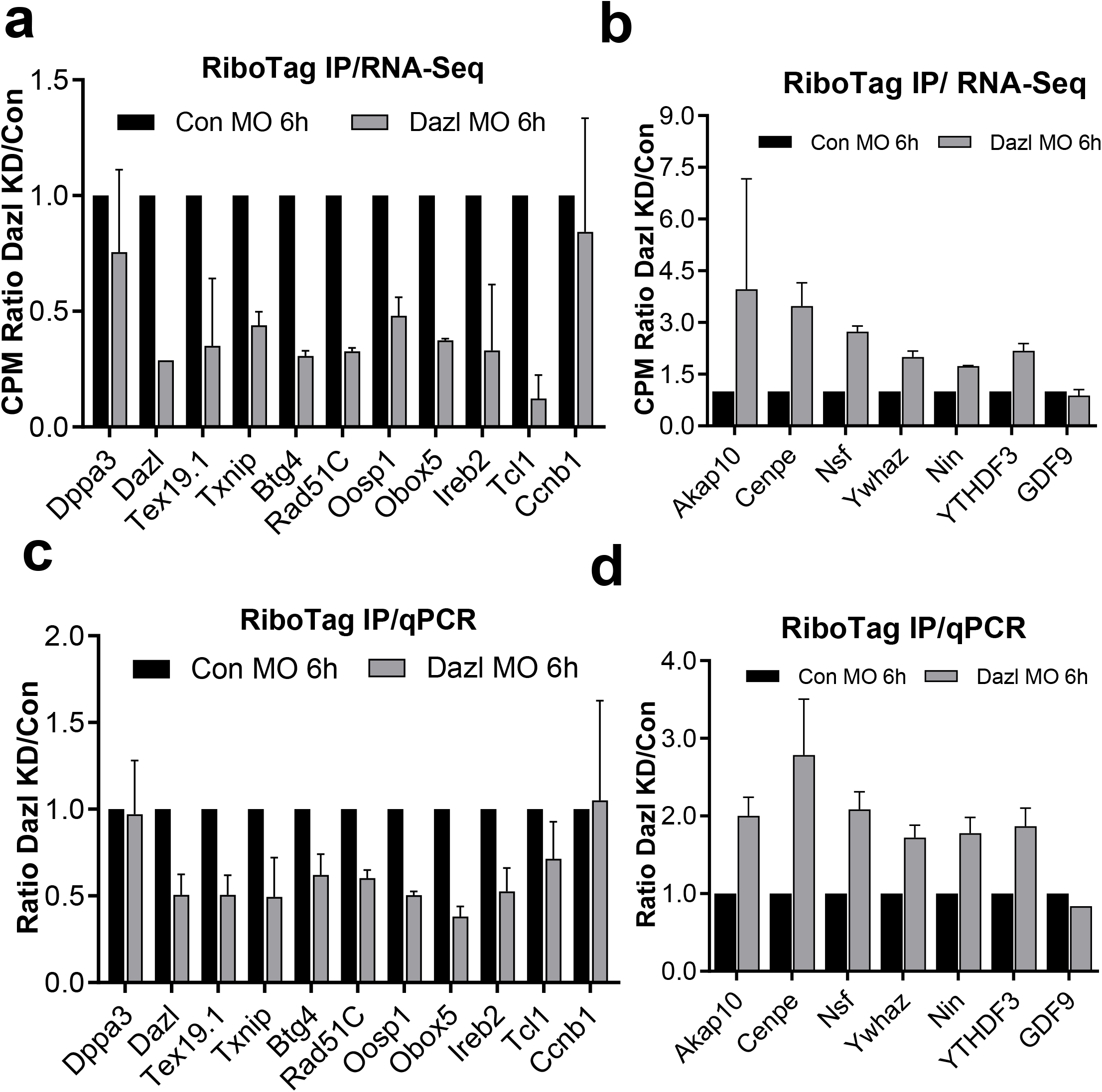
RiboTag IP/qPCR confirms the presences of a subset of transcripts whose translation is upregulated and downregulated in oocytes depleted of DAZL. Representative targets affected by DAZL removal in RiboTag/RNA-Seq dataset are showed in panel **(a)** (transcripts whose translation is downregulated by DAZL removal) and **c** (transcripts whose translation is upegulated by DAZL removal); the differences in ribosome loading DAZL MO/CON-MO for the same transcripts assessed in independent biological replicates by RiboTag IP/qPCR is reported in panel **(b)** and **(d)**. *Dppa3* and *Ccnb1* are used as negative control in panel **(a)** and **(b)** for the transcripts whose translation is downregulated by DAZL removal, as they are not regulated by DAZL during oocyte maturation. *Gdf9* mRNA is used as negative control in panel **(c)** and **(d)** for the transcripts whose translation is upregulated by DAZL removal, as it is not regulated by DAZL during oocyte maturation. Wild type and *Dazl*^+/−^ mice were injected with control or DAZL-MO. After overnight preincubation with 2 μM milrinone, oocytes were cultured in inhibitor-free medium for maturation. Oocytes were collected at 6 hrs for RiboTag IP and qPCR analysis.

### DAZL physically interacts with transcripts whose translation is upregulated or downregulated after DAZL removal

The above findings open the possibility that DAZL binding to maternal mRNA leads to both increase and decrease in translation. If these were correct, DAZL should bind to maternal mRNAs whose translation is upregulated or downregulated during oocyte maturation. To test this hypothesis, we analyzed a previously generated DAZL RIP-Chip dataset. In this DAZL RIP-Chip dataset, 811 transcripts are significantly immunoprecipitated (> 1.5 fold enrichment as compared to IgG) by DAZL antibodies during the GV to MII transition. A scatter plot (Fig. 4a) of these data shows DAZL binding to both upregulated transcripts and downregulated transcripts, a finding consistent with the two classes distribution of the ribosome loading transcripts (Supplementary Fig 2 a). This analysis is again consistent with the hypothesis that DAZL interacts with both classes of transcripts whose translation may increase or decrease during oocyte maturation. A more quantitative comparison of the mRNA immunoprecipitated by DAZL antibody and the transcripts whose translation is affected by DAZL depletion shows that 212 downregulated and 49 upregulated (total 251) transcripts are also immunoprecipitted by DAZL antibodies. A sizable number of transcripts (215) are not affected or changes do not reach statistical significance (Supplementary Fig. 4).

**Figure 4.**
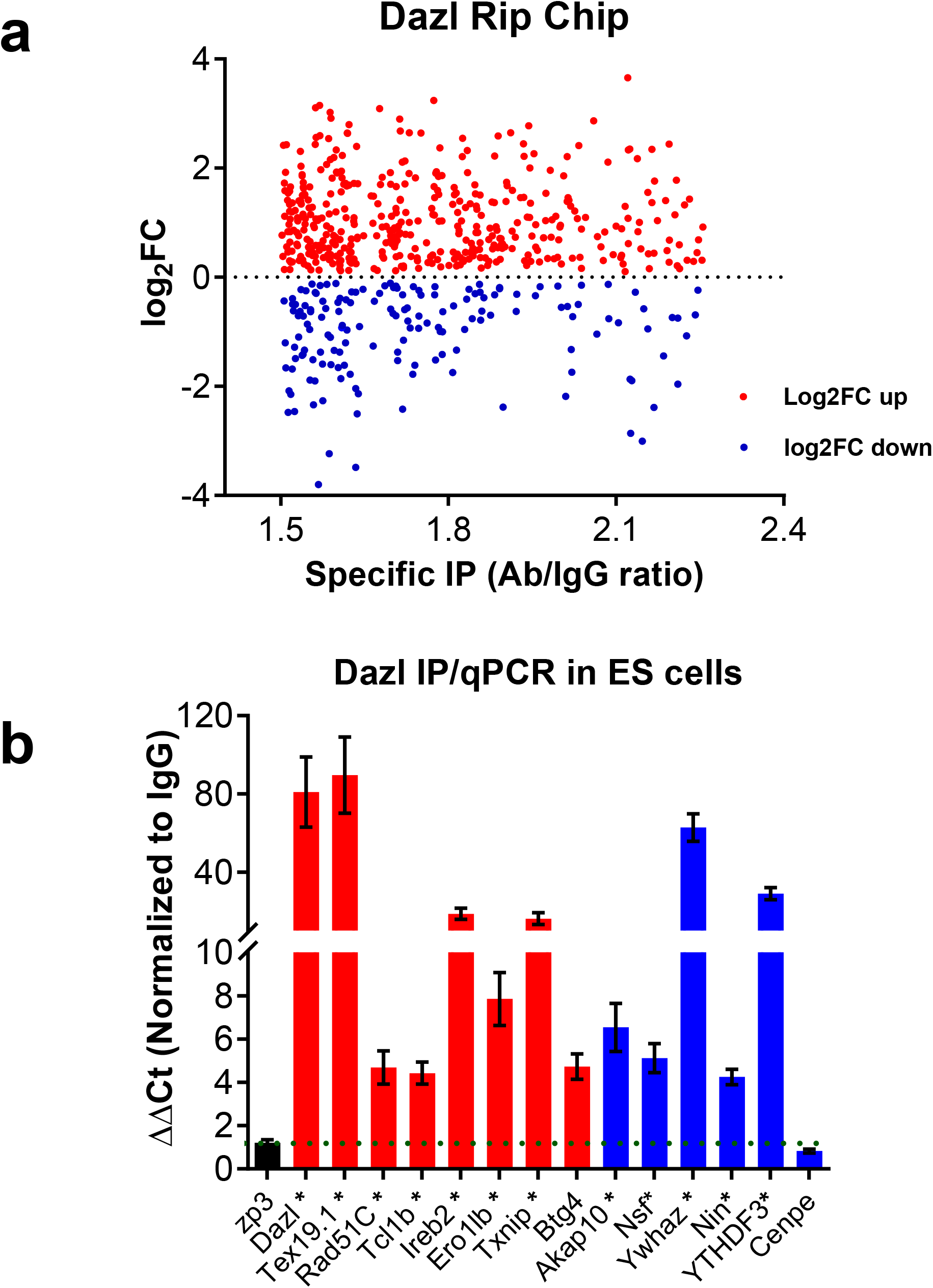
DAZL interacts with transcripts whose translation is upregulated or downregulated during oocyte maturation. **a.** Comparison DAZL TipChip/RiboTag IP RNseq in oocytes. Changes in ribosome loading from 0 hr (GV) to 16 hrs (MII stage) of DAZL targets assessed by RIP-Chip. A subset of transcripts whose translation increased from GV to MII stage are also enriched in DAZL immunoprecipitates of oocyte extracts (red); transcripts whose translation decreases during oocyte maturation are specifically immunoprecipiated by DAZL Antibody (blue). wild type oocytes were collected at 0h and 6hrs of *in vitro* maturation for RiboTag IP/RNASeq analysis. For DAZL RIP-Chip, wild type oocytes were primed with PMSG and after stimulation with hCG, MII stage oocytes were harvested as described. Oocyte lysates was immunoprecipited with DAZL-specific antibody or IgG and the mRNA recovered in the IP pellet measured by microarray hybridization. (RIP-Chip data kindly provided by Jing Chen and Mat Cook members of the lab) (b) DAZL RNA-IP qPCR of ES cells extracts. ES cells were cultured in DMEM medium with supplements include 15% KOSR, 2% FBS, 1x MEM Non-Essential Amino Acids (100x), 1x 2-mercaptoethanol, 10^-7^ U/ml LIF, 1x Pen/Strep and 2i (1 μM PD0325901 and 3μM CHIR99021) and collected for DAZL RNA-IP/qPCR analysis. The results are normalized with IgG IP. Asterisks (*) indicate that these transcripts are also found to be associated with DAZL also in the RIP-Chip dataset.

We wished to next confirm that transcripts whose translation increases after DAZL depletion are indeed direct target of DAZL. Because no sufficient signal could be obtained in DAZL RIP when using 200 oocytes with currently available antibodies, we used mouse embryonic stem cells(ES) for DAZL RIP. It is established that ES cells in the ground state express DAZL as well as a large number of two-cell embryo transcripts, mRNAs often expressed also in oocytes^15, 22 23^. ES cells were cultured in DMEM medium +2i and collected for DAZL IP/qPCR analysis. The results are normalized to the IgG IP. This DAZL RIP experiment in ES cells shows that transcripts of *Dazl, Tex19.1, Txnip, Rad51C, Btg4, Ero1lb, Ireb2*, and *Tcl1b* (Fig. 4B, red), whose translation is downregulated by DAZL removal, and transcripts of *Akap10, Nsf, Ywhaz, Nin*, and *YTHDF3*, whose translation is upregulated by DAZL removal (Fig. 4b, blue), are all specifically immunoprecipitated by DAZL antibody (Fig. 4b). However, *Cenpe*, a transcript whose translation is upregulated by DAZL removal, could not be immunoprecipitated by the DAZL antibody, suggesting that DAZL may also act as a translational repressor through an indirect pathway. Nevertheless, most of the transcripts whose translation increases/decreases after DAZL depletion are directly interacting with DAZL in oocytes and ES cells.

### The 3’ UTR of representative transcripts Oosp1 and Obox5 recapitulates the effect of DAZL depletion on translational activation

*Oosp1* and *Obox5* are two oocyte-specific transcripts whose translation is affected by DAZL removal as determined in both the RiboTag IP/RNA-Seq dataset and in the RiboTag IP/qPCR validation. OOSP1 (oocyte secreted protein 1) was initially identified as a novel oocyte-secreted protein ^24^. OBOX5 (oocyte specific homeobox 5) is a member of the Obox family of proteins but its function is unclear ^25^. These mRNAs were chosen because of their robust translational activation in meiosis and their relatively simple 3’UTR. To verify the effect of DAZL on translation of these two candidate mRNAs, a YFP reporter was fused to the *Oosp1* or *Obox5* 3’UTR and these constructs were injected in oocytes together with either Con MO or DAZL MO. A fully polyadenylated *mCherry* reporter was used as a control of the volume injected. The accumulation of YFP and mCherry in individual oocytes was recorded throughout meiotic maturation and YFP signals were corrected by the level of co-injected mCherry signal. Data are expressed as changes over 0 hr (GV stage), as differences in reporter accumulation were detected in GV-arrested oocytes (see below). By measuring the average YFP signals throughout maturation, the accumulation of *YFP-Oosp1* and *YFP-Obox5* reporter in CON-MO group closely follows the corresponding ribosome loading onto the endogenous mRNA; however, DAZL depletion causes at least 50% decrease in translation in Oosp1 and Obox5 reporter during oocyte maturaion (Fig. 5a, c). We further assessed the rates of translation during oocyte maturation of the two reporters by fitting the YFP/mCherry data during GV (0-2 hrs) and after GVBD (4-8 hrs) (Fig. 5b, d). We found a significantly decrease of Oosp1 (p<0.0001) and Obox5 (p<0.0001) translation rates in DAZL MO injected oocytes (Fig. 5b, d), confirming that DAZL depletion decreases the translation of these reporters during oocyte maturation. Consistent with our RiboTag IP/RNASeq data and translational efficiency of *Oosp1* and *Obox5* is affected by DAZL depletion (Supplementary Fig. 5). *Ccnb1* 3’UTR co-injected with either CON-MO or DAZL-MO shows no obvious changes in translational accumulation between the two groups, confirming the selective effect of the DAZL depletion (Fig. 5e, f).

**Figure 5.**
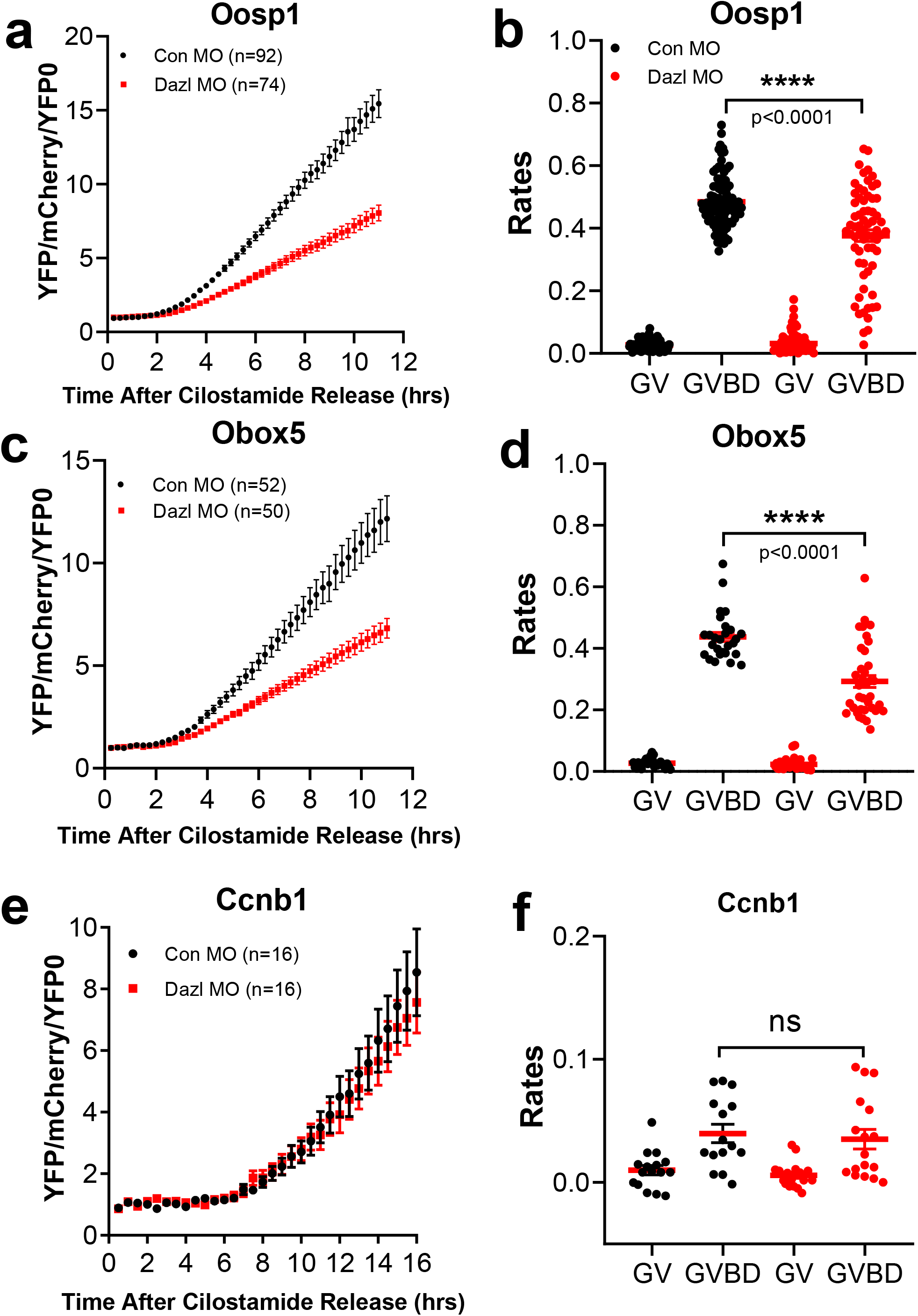
The 3’ UTR of Oosp1 and Obox5 recapitulates the effect of DAZL depletion on mRNA translation. Oocytes were injected with 12.5ng/uL *mCherry-polyadenylated* mRNA and 12.5ng/uL *YFP-Oosp1 3’UTR* reporter for or *YFP-Obox5 3’UTR* reporter with either CON-MO or DAZL-MO. Oocytes were then pre-incubated overnight to allow the mCherry signal to reach a plateau. At the end of the preincubation, oocytes were released in cilostamide-free medium for maturation and YFP and mCherry signal were recorded by time lapse microscopy at a sampling rate of 30 min for 12 hrs. The YFP signal were corrected by the level of coinjected mCherry signal and normalized to the first reconrding of YFP/mCherry. Experiments were repeated three times and the data are the cumulative mean±SEM of three independent experiments. DAZL depletion decrease translation of reporter driven by the *Oosp1* **(a)** or *Obox5* **(c)** 3’UTR during oocyte maturation, while *YFP-CcnB1 3’UTR* **(e)** is not affected. Individual oocyte YFP/mCherry were used to calculate the rate of translation of the reporters at the 0-2 hrs (prior to GVBD) and 5-10 hrs (after GVBD). The rates of *YFP-Oosp1* **(b)** (p < 0.0001) or *YFP-Obox5* **(d)** (p < 0.0001) translational accumulation are significantly decreased in GVBD after DAZL removal, whereas the rates of *YFP-CcnB1* **(f)** translation are not significantly changed.

To confirm that the depletion of DAZL protein with the specific MO is the sole cause of the decreased translation of the reporter, we performed the following rescue experiment. A human recombinant DAZL protein was injected together with the DAZL MO and the Oosp1 reporter. As observed above, DAZL depletion causes a decrease in the rate of translation of the Oosp1 reporter. This decrease is completely rescued when the recombinant DAZL protein is coinjected with the DAZL MO (Fig. 6a, b). The rescue effect of the DAZL protein was not limited to the translation efficiency. As previously reported, DAZL depletion on a het background almost completely prevents oocyte maturation to MII (Control MO 69.7 % vs DAZL MO:7%). Conversely, when the DAZL MO is co-injected with a recombinant DAZL protein, oocytes complete maturation to MII at a rate (63%) similar to control injected oocytes(Fig. 6c). Taken together with the RiboTag IP / RNA-Seq and qPCR data (Fig. 2, Fig. 3), the reporter measurements further support the conclusion that the DAZL protein plays a role in the translational activation of these two target mRNAs, that their 3’UTR mediates the effect of this RBP on translation, and that DAZL depletion is the cause of decreased translation.

**Figure 6.**
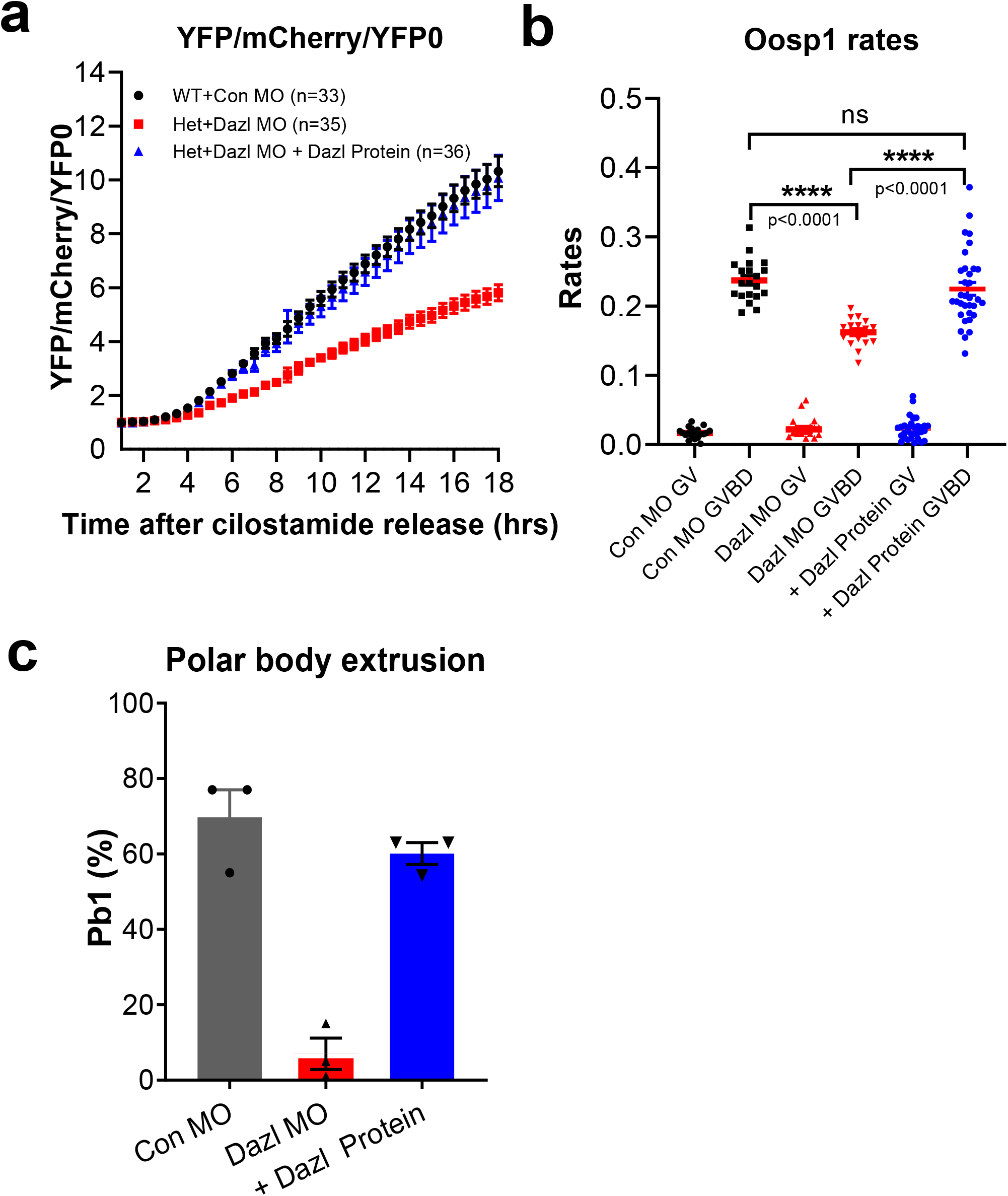
The translation of the YFP-Oosp1 reporter is rescued by DAZL protein. **(a)** Human DAZL protein injection restores Oosp1 translation during oocyte maturation in MO injected oocytes. Oocytes were injected with 12.5ng/uL *mCherry* mRNA and12.5ng/uL *YFP-Oosp1 3’ UTR* reporter with either CON-MO or DAZL-MO with or without recombinant human DAZL protein, and incubated in cilostamide medium overnight to allow mCherry signal to reach a plateau. At the end of the preincubation, oocytes were released in cilostamide-free medium for maturation and YFP and mCherry signal recorded by time lapse microscopy at a sampling rate of 30 mins for 12 hrs. YFP signal were corrected by the level of co-injected mCherry signal and were normalized to the first time point. Experiments were repeated 3 times and the data are the mean + SEM of three independent experiments. **(b)** Human DAZL protein injection restores the rate of *YFP-Oosp1* translation with the effect of DAZL depletion in GVBD to levels of CON-MO. **(c)** Microinjection of a human DAZL protein rescues the meiotic block of oocytes injected with DAZL-MO. Oocytes maturation was scored by counting the number of oocytes with a polar body. The injection of this human DAZL protein restored polar body formation to levels not significantly different from control (63% versus 70%). Three independent experiments were performed and the data reported here are the average ratio of polar body extrusion.

### DAZL depletion increases the translation of Oosp1 and Obox5 mRNAs in GV-arrested oocytes

In the above experiments on reporter translation, we consistently observed that translation of both reporters during the first two hours of incubation, when the oocytes are still GV-arrested, is significantly increased in the DAZL-MO injected group (Oosp1: p < 0.0001 and Obox5 p = 0.0007) (Fig. 7b, d). To verify this apparent de-repression in DAZL-depleted, quiescent oocytes, we reanalyzed the RiboTag IP/RNASeq data sets. We found that ribosome loading on both Obox5 and *Oosp1* during GV-arrested is also increased in this dataset (Supplementary Fig. 6). To remove any possible bias due to variation in the total mRNA, we next calculated the translational efficiency (TE) of these two transcripts after DAZL depletion. Indeed, the TEs for *Oosp1* and *Obox5* are significantly increased in GV oocytes depleted of DAZL (Fig. 7a, b), whereas the *CcnB1* translation is not affected (Fig. 7e) To assess whether this effect of DAZL depletion is widspread, we reanalyzed the RiboTag IP RNASeq data and found that ribosome loading of approximately 70 transcripts is significantly increased after DAZL removal in GV oocytes (69, Supplementary Fig. 6 red, FDR<0.05). This latter finding provides further evidence for a repressive role of DAZL prior to meiotic resumption.

**Figure 7.**
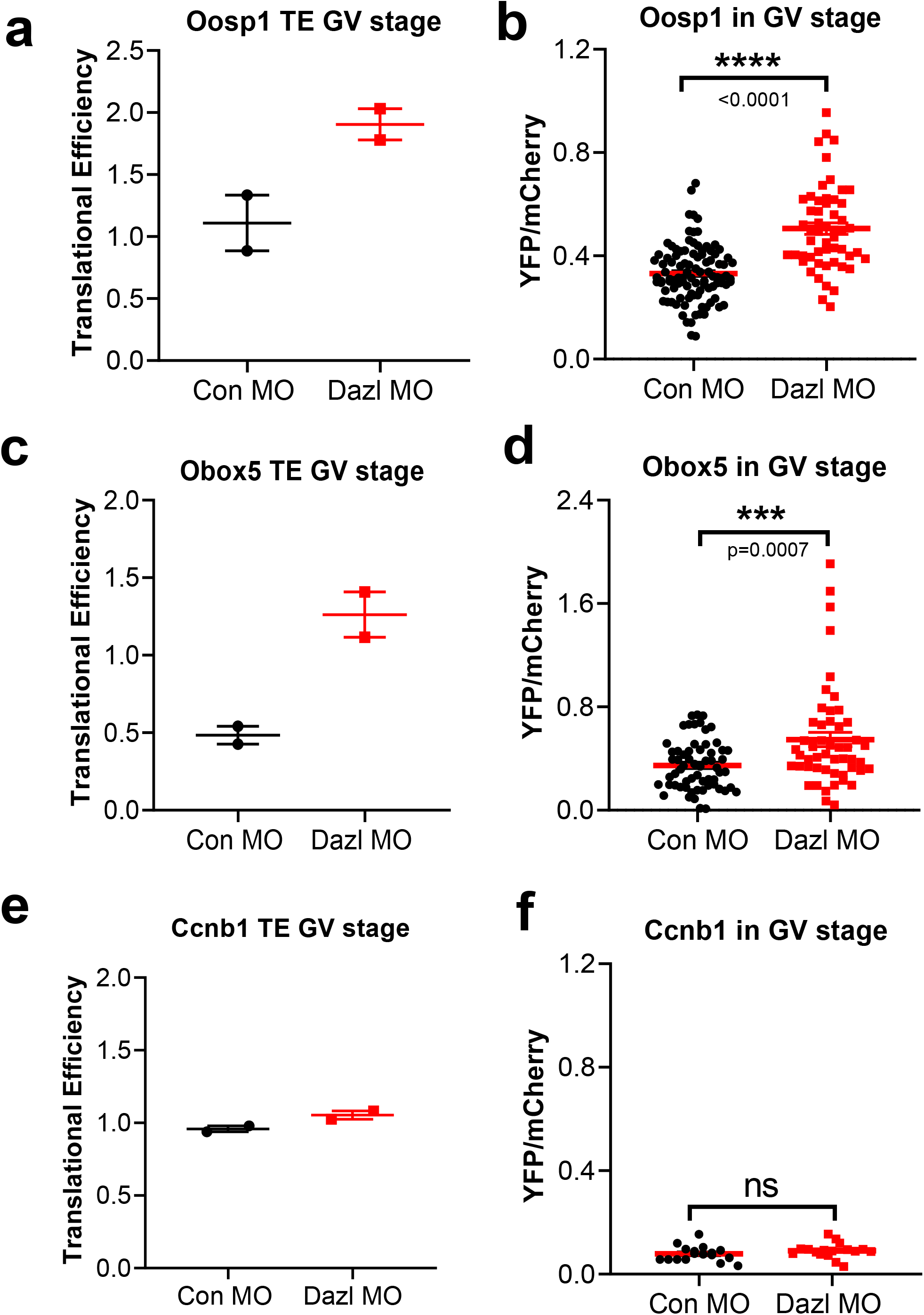
DAZL depletion increases translation of Oosp1 and Obox5 endogenous transcripts and Oosp1 and Obox5 reporters in GV-arrested oocytes. **(a, c, e)** GV stage oocytes from wild type or *Dazl*^+/−^ mice were injected with CON-MO or DAZL-MO. Oocytes were preincubated overnight with 2 μM milrinone and then cultured in inhibitor-free medium for maturation. 0hr (GV stage) data from RiboTag IP/ RNA-Seq was used for Tra nslational efficiency (TE) analysis. TE was calculated as the ratio of the CPM for HA immunoprecipitated transcripts *Oosp1* or *Obox5* over the corresponding input at 0 hr oocyte. The TEs for *Oosp1* **(a)** and *Obox5* **(c)** is increased in GV oocytes depleted of DAZL. However, TE for *CcnB1* **(e)** is not affected. **(b, d, f)** GV stage oocytes were injected with 12.5ng/uL *mCherry-* polyadenylated mRNA and 12.5ng/uL *YFP-3’ UTR* reporter for *Oosp1* 3’ UTR or *Obox5* 3’UTR with either COMN-MO or DAZL-MO. Oocytes were pre-incubated overnight to allow mCherry signal to plateau, then released in cilostamide-free medium. YFP signal were corrected by the level of coinjected mCherry signal. The translation of both reporters in GV-arrested oocytes is significantly increased *Oosp1* (p < 0.0001) (b) and *Obox5* (p = 0.0007) **(d)** in DAZL-MO injection group, whereas no significant difference in the translation of the *CcnB1* reporter (f) is detected. Experiments were repeated three times and the data reported are the rates for each individual oocytes from three independent biological replicates.

### Efficient translation of Oosp1 and Obox5 reporters requires the presence of DAZL binding element

Collectively, the above experiments strongly indicate a translational regulation of the *Oosp1* and *Obox5* transcripts are affected by DAZL MO injection during GV to MI transition. To test whether this effect is due to binding of DAZL to these target mRNAs, we mutated the consensus putative DAZL-binding sites (UU[G/C]UU) by replacing critical nucleotides with adenosine in *Oosp1* 3’UTR or *Obox5* 3’UTR. We have previously shown that this mutations disrupts DAZL binding ^19, 26^. A schematic representation of *Oosp1* and *Obox5* 3’UTR with mutant DAZL binding sequence is reported in Fig. 8a. When YFP reporter tagged with mutant *Oosp1* 3’UTR or *Obox5* 3’UTR YFP reporters were injected in oocytes and their rate of translation were monitored during maturation, mutation of a single DAZL-binding site in either *Oosp1* or *Obox5* 3’UTR is sufficient to significantly decrease the rate of reporter accumulation (Fig. 8c, d, f, g) during meiotic resumption. Of note, mutation of DAZL binding site of *Oosp1* (Fig. 8b) or *Obox5* (Fig. 8e) also cause an increase translation in GV stage oocyte as compared to wildtype reporters. This increased rate of translation of the reporter was confirmed in a different experimental paradigm where control and DAZL depleted oocytes are maintained in GV stage. Also under these conditions, the translation rate of the *Oosp1* reporter is increased, whereas the *CcnB1* mRNA is not affected (Supplementary Fig. 7) This latter finding is consistent with the results of DAZL MO of the RiboTag IP/RNASeq experiment (Fig. 7a, c), confirming that DAZL functions as translational repressor also in GV-arrested oocyte.

**Figure 8.**
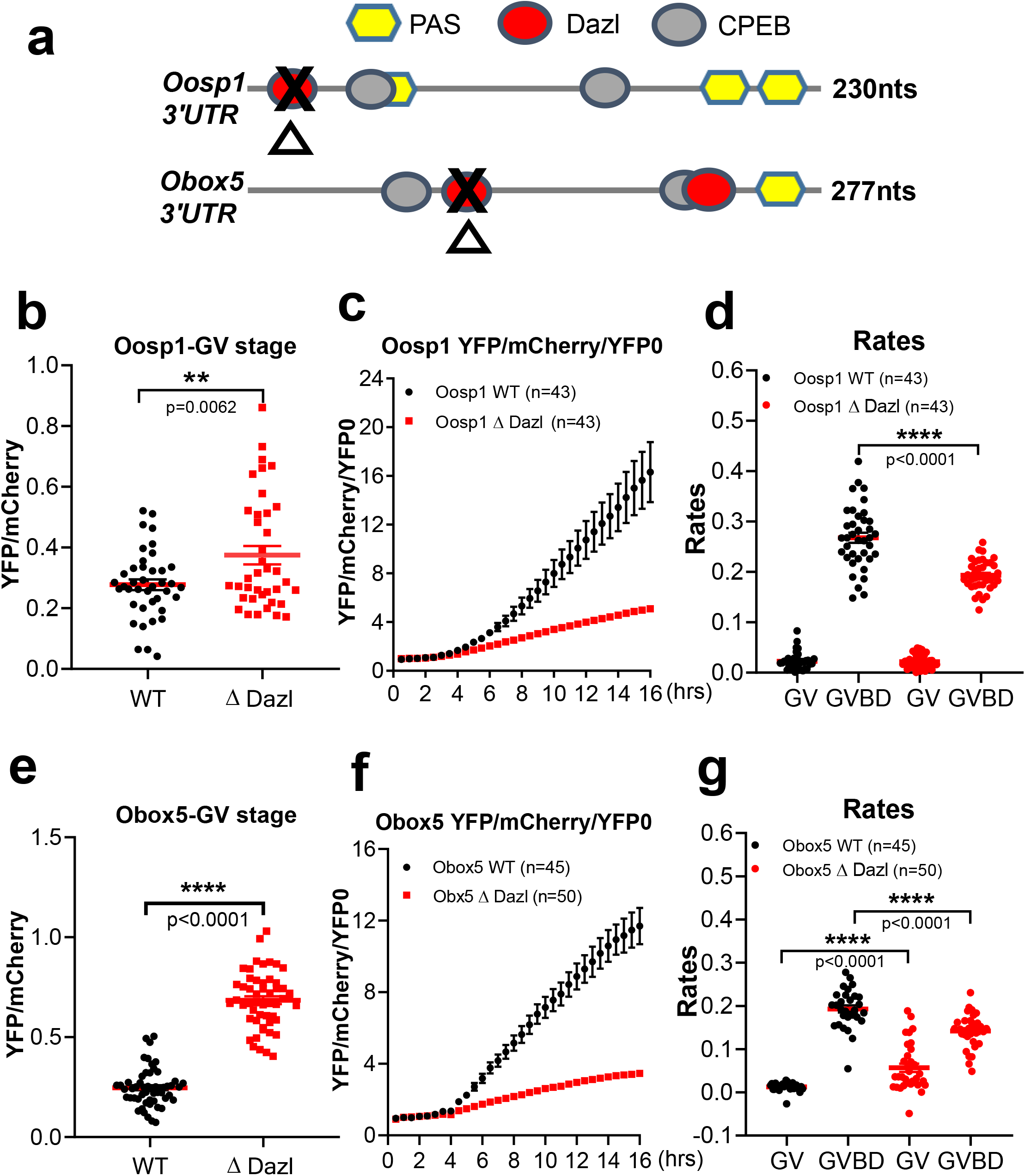
Translation of Oosp1 and Obox5 reporter is dependent on the presence of a DAZL binding element. Constructs with the mutated DAZL-binding sequence, with wild type *Oosp1 3’ UTR*, or with the *Obox5 3’UTR* along with mCherry-polyadenylated mRNA were injected into GV stage oocytes at 12.5ng/uL per reporter. After overnight pre-incubation to allow mCherry signal to plateau, oocytes were released in cilostamide-free medium and recorded under the microscope for tracking *YFP-Oosp1* or *YFP-Obox5* translation during oocyte maturation. YFP and mCherry images were acquired every 30 mins for 18 hrs. YFP signal were corrected by the level of co-injected mCherry signal. Experiments were repeated three times on different days. **(a)** Scheme of the *Oosp1* and *Obox5* 3’ UTR and position of the PAS, and putative CPEB and DAZL-binding elements. Mutagenesis of the putative DAZL-binding element was performed as detailed in ‘Materials and Methods’ sectioni. A red oval is the DAZL consensus sequence in the 3’UTR of *Oosp1* and *Obox5*. A black cross indicates the mutated DAZL-binding consensus sequence. **(b and e)** The effect of DAZL-binding element mutation on *Oosp1* (p = 0.0062) or *Obox5* (p < 0.0001) translation in GV stage. Rates of reporter accumulation were calculated in each GV-arrested oocyte and plotted as individual dots. Mean and SEM values were calculated from all the oocytes measured from three different experiments. **(c and f)** Mutation of DAZL-binding element on 3’UTR of *Oosp1* or *Obox5* decreases translation of each respective reporter during oocyte maturation. YFP signals were corrected by the level of co-injected mCherry signal. Every time point was normalized to the first time point of YFP:mCherry. **(d and g)** The rates of *YFP-Oosp1* (p<0.0001) or *YFP-Obox5* (p < 0.0001) reporter accumulation during maturation is significantly decreased in GVBD after mutation of the putative DAZL site.

## Discussion

With the experiments described above, we provide evidence that the RNA-binding protein DAZL plays a function in fully-grown oocytes by shaping the pattern of maternal mRNA translation at this critical transition of gametogenesis. Our data document that DAZL has both inhibitory, and stimulatory, effects on translation in quiescent oocytes as well as during meiotic resumption. This dual function parallels that of CPEB1, which is considered to be the master regulator of translation in oocytes, and reinforces the concept that DAZL cooperates with CPEB to repress and active translation of maternal mRNAs. Given the finding that DAZL-MO treatment prevents progression through meiosis, it is proposed that the DAZL function is essential for oocyte maturation.

Several lines of evidence support the conclusion that DAZL is expressed and functional at the end of oogenesis in fully grown mouse oocytes. The mRNA coding for DAZL is expressed at high levels in fully-grown MII oocytes^18^. RiboTag IP/RNASeq data confirmed by qPCR document that the *Dazl* mRNA is actively translated and translation increases during oocyte maturation. In line with mRNA ribosome loading, DAZL protein is detected by Western blot with two distinct antibodies. The progressive increase in the DAZL protein during maturation is consistent with the increase in translation, further strengthening the view that this RBP accumulates during oocyte maturation. Finally, the RIP-Chip data document that the DAZL is actively interacting with hundreds of maternal mRNAs. All these independent pieces of evidence strongly support the hypothesis of expression and function if this RBP at the final stage of oogenesis.

The function of DAZL in translation is further confirmed by a loss-of-function approach. Morpholino oligonucleotide interference with mRNA translation was used on a DAZL-heterozygous background, since homozygous deletion of this gene precludes oocyte development. As detailed above, we chose to determine the effect of DAZL effect on translation in MI to avoid secondary effects due to the block in oocyte maturation. Injection of DAZL-MO interrupts *Dazl* mRNA translation as detected by RiboTag IP/qPCR. In parallel with decreased translation, Wester blot analysis of oocyte extracts shows a reproducible decrease of more than 90% in DAZL protein. The specificity of this treatment is supported by the data showing that ribosome loading onto the mRNAs for *CcnB1, Dppa3* and *Gdf9* is not affected. At the transcriptome level, the overnight incubation to deplete the DAZL-MO injected oocytes has minimal effect on total transcript levels, arguing against a generalized disruption of the oocyte viability. All these findings increase confidence that the KD of DAZL is effective in depleting the oocytes of the DAZL protein and that its effect is specific.

The analysis of the translatome in DAZL-depleted oocytes indicates that approximately 800 maternal mRNAs show altered on the level of ribosome loading. Together with the DAZL RIP-Chip data, this analysis confirms the presence of a large number of maternal mRNA targets for this RBP. *Tex19.1, Txnip, Rad51C, Btg4, Oosp1, Obox5, Ireb2, andTcl1* are examples of the more than 200 mRNAs regulated by DAZL on the basis of the decreased ribosome loading after DAZL depletion and the observation that these mRNAs are immunoprecipitated by DAZL antibodies. These findings confirm our and others previous observations that *Tex19.1* mRNA is a DAZL target^19, 27^. TEX19.1 protein accumulates during oocyte maturation and data in the testis indicate that this protein may be involved in the regulation of transposon expression^28, 29, 30^. Similarly, DAZL regulation of *Btg4* mRNA translation has been reported by others^31, 32, 33^. In both mouse and human, *Btg4* transcripts are highly enriched in the ovary and testis. A consequence of the absence of BTG4 is a global delay in maternal mRNA degradation during the MZT^31, 32, 33, 34^. Given the involvement of BTG4 in mRNA destabilization, one would expect that DAZL depletion would induce mRNA stabilization by preventing Btg4 accumulation. However in our experimental paradigm, DAZL depletion and consequent decreases in BTG4 accumulation at 6 hrs are probably not sufficient to produce detectable effect on mRNA stabilization. mRNA destabilization is detected at 8 hrs of oocyte maturation (data not shown). Finally, it should be noted that ribosome loading onto all the above listed mRNAs increases during oocyte maturation and most of the mRNAs also contain at least one putative CPE element in the 3’UTR. Thus, one function of DAZL is likely to increase the translation of these mRNA during maturation, a function likely synergistic with that of CPEB, as we have described for *Tex19.1* ^19^. This conclusion is supported by the reporter translation of *YFP-Oosp1* and *YFP-Obox5*. Similarly, the data with the *YFP-CenpE* reporter (Supplementary Fig. 8) are consistent with the view that a group of transcripts, that include *Akap10, Cenpe, Nsf, Ywhaz, Nin*, and *YTHDF3*, are translationally repressed by DAZL.

Our DAZL RIP-Chip data indicates that DAZL interacts with approximately 800 mRNAs expressed in the oocytes. The interaction of several candidate mRNAs with DAZL was also confirmed in ES cell extracts. Data are available for *Dazl* mRNA targets in the fetal gonad of the mouse and human. Of the mRNA shown to interact with DAZL in human fetal ovary^35^, *Sycp1* and *Tex11* are not detected in GV oocytes and the ribosome loading of *Smc1b* although decreased in our data set does not reach statistical significance. A comparison between the human fetal gonad data ^35^ and our mouse RIP data shows overlap in immunoprecipitation of 72 mRNAs including *Trip13, Rad 51, Spin1, Kit*, and *Arpp19*.

One consistent finding is that *Dazl* mRNA is immunoprecipitated with DAZL antibody in both mouse and human fetal ovary. This finding reinforces the idea that DAZL is involved in an aoutregulatory loop controlling its own translation^18^.

Of the mRNAs identified in the DAZL RIP-Chip, 261 of 476 transcripts are affected by DAZL depletion (Supplementary Fig. 4). The limited overlap between the transcripts recombined in the RIP and in the RiboTag IP/RNASeq is in part due to the differences in annotation of the two platforms (300 genes not shared) or to the different filtering of the data. Our numbers are also considerably lower than those reported by Zagore et al. using HITS-CLIP with testis extracts ^17^. This latter difference may be due to the fact that we did not use crosslinking for our experiment but also to the fact that very low amounts of cell protein is available from the oocyte. Thus, the affinity of the antibody for DAZL becomes limiting. Similar to the finding of Zagore et al.^17^, we find that about 200 mRNAs that bind DAZL are not affected by DAZL depletion. This discrepancy again can be due to filtering of the data and the cutoffs imposed. Also, we should point out that in our experimental paradigm, we are measuring acute effects of DAZL depletion and if longer incubation times are used the number of mRNAs affected would increase substantially.

A previous report in the testis proposes that DAZL is involved in mRNA stabilization^17, 36, 37, 38^. Therefore it is possible that DAZL depletion affects translation indirectly by destabilizing mRNAs. However, overnight depletion of DAZL has minimal effects on the oocyte transcriptome, lessening the possibility of destabilizing effects on maternal mRNAs. Moreover, all the data of the candidates we more thoroughly investigated are inconsistent with the destabilization hypothesis, as we cannot detect a decrease in total mRNA. Therefore, the decreased translation for these candidate DAZL targets is not due to destabilization of these mRNAs. Moreover, many of the effects of DAZL depletion on translation continue to be present when the TE is calculated, indicating that mRNA stabilization is not a factor in ribosome loading. However, the timeframe of our experiments is considerably short and we cannot exclude that longer time causes of DAZL depletion uncovers effects of DAZL depletion on mRNA stability.

Recently, it has been reported that DAZL is dispensable for oocyte maturation, but that instead its overexpression has deleterious effects on oocyte developmental competence^20^. This conclusion is based on the observation that DAZL protein is markedly decreased in adult ovary in comparison with neonatal ovary; however, the variable ratio somatic:germ cells in the gonad during development may account, at least in part, for these differences^20^. Our data on DAZL protein expression detected with two distinct antibodies, the RIP-Chip data, and the translational regulations described confirm the expression and increased accumulation of DAZL in the final stages of oocyte development. Genetic manipulations also led to the conclusion that the absence of DAZL does not produce overt phenotypes on oocyte maturation or fertility. The genetic background used in these experiments is a mixed background (ICR and C57BL/6N) while we use a pure C57 BL/6 background. It has been noted that the penetrance of the phenotypes associated with *Dazl* gene ablation are sensitive to the mouse background used^12^. However, the view that a DAZL needs to be expressed within a very narrow range of concentrations is consistent with our findings that DAZL has a dual effect on translation, functioning both as a repressor and activator. Therefore, it is possible that increased DAZL levels favors translational repression that would be detrimental to developmental competence. Aside from the genetic background of the mice used, not immediately evident is the explanation of why Dazl KO in neonatal oocytes produces no detectable phenotype on fertility. Possible off-target effects of DAZL morpholinos are inconsistent with the rescue experiments we have performed. In several cases, it has been observed that morpholino oligonucleotide treatment is associated with induction of *p53* ^39, 40^(*tp53* or *trp53*) or interferon response or toll like receptor^41^. Since transcription is repressed in GV oocytes, it is unlikely that MO off-target effects include changes in transcription. However, we noticed that *trp53* mRNA becomes associated with ribosome during oocyte maturation and is immunoprecipitated with DAZL antibody. We could not detect clear effects of DAZL depletion on *trp53* mRNA translation. Another possible explanation of the divergent observations is that the oocyte does not tolerate acute depletion of DAZL, while it has time to adjust to loss of DAZL during the follicle growth phase that starts in the neonate ovary. Genetic compensation has been shown to be at the basis of differences in phenotypes produced by mutations but not knockdowns^42^. Since we have shown that DAZL functions in partnership with CPEB^19^, it is possible that this latter RBP would compensate for the loss of DAZL. In this respect, it would be important to determine whether CPEB expression is affected in the DAZL KO and how the translational program is executed in the absence of DAZL in the fully grown oocyte.

Several scenarios may explain the dual repression/activation role of DAZL on translation. One possibility is that DAZL assembles different molecular complexes in GV-arrested and MI oocytes. DAZL has been shown to interact with PUM2 forming a translational repressor complex on the *Ringo/Spy* mRNA^43^. During maturation, the complex is dissociated leading to translational activation. Thus, one could envisage that DAZL is part of a repressive complex in mouse GV-arrested oocytes and contributes to an activating complex in MI stage oocytes. This scenario is reminiscent of the CPEB1 mode of action^8, 44^. We have observed that the DAZL protein shifts in mobility on SDS/PAGE during oocyte maturation, suggesting that the protein becomes post-translationally modified during maturation. Finally, it should be considered that the concentration of DAZL protein increases up to six fold during oocyte maturation. Thus, it may be possible that low loading on an mRNA is sufficient to repress translation, whereas loading of multiple DAZL proteins on a mRNA leads to activation of translation. Indeed, we and others have proposed that the number of loaded DAZL synergizes in activation of translation^14, 18^.

In summary, our findings are consistent with a role of DAZL in the translation program executed during the final stages of oocyte maturation. The dual function as repressor and activator suggests that complex changes in the proteome in fully grown oocytes and during maturation are dependent on DAZL action. These findings imply that spontaneous DAZL mutations found in human may affect not only germ cell development in the fetal gonad but they also have an effect on oocyte quality. Such a possibility has been proposed with the description of missense mutations in infertile women^45^.

## Materials and Methods

### Experimental animals

All experimental procedures involving animal models used were approved by the Institutional Animal Care and Use Committee of the University of California at San Francisco (protocol AN101432). Pure C57BL/6 female mice (21-24 days old) carrying the DAZL TM1^Hgu^ allele (Δdazl) were generated as previously described^18, 19^. Rpl22tm1.1Psam/J (RiboTag) mice, with a targeted mutation that provides conditional expression of the ribosomal protein L22 tagged with three copies of the HA epitope. Rpl22tm1.1Psam/J homozygous males were crossed with C57BL/6-TgN (Zp3-cre) 82Knw (Jackson Laboratories) females to produce C57BL/6-Zp3cre-Rpl22tm1.1Psam (Zp3cre-RiboTag) mice. For breeding Zp3RiboTagDazl^+/+^ or Zp3RiboTagDazl^+/−^, C57BL/6-Zp3cre-RiboTag wild type or homozygous males were crossed with C57BL/6-RiboTag wild type or heterozygous ΔDAZL females to obtain C57BL/6-ΔDAZL-ZP3cre-RiboTag mice.

### Oocyte collection and microinjection

Oocyte isolation and microinjection were performed using HEPES modified minimum essential medium Eagle (Sigma-Aldrich, M2645) supplemented with 0.23 mM pyruvate, 75 μg/mL penicillin, 10 μg/mL streptomycin sulfate, and 3mg/mL BSA, and buffered with 26 mM sodium bicarbonate. To prevent meiosis resumption, 2 μM cilostamide (Calbiochem, 231085) was added in the isolation medium. Oocyte *in vitro* maturation was performed using Eagle’s minimum essential medium with Earle’s salts (Gibco, 12561-056) supplemented with 0.23 mM sodium pyruvate, 1% streptomycin sulfate and penicillin, and 3mg/mL bovine serum albumin (BSA). For microinjection, cumulus cells were removed by mouth pipette from isolated cumulus oocyte complexes (COCs) and denuded oocyte were injected with 5-10 pL of 12.5 ng/μL mRNA reporter using a FemtoJet Express programmable microinjector with an Automated Inverted Microscope System (Leica, DM4000B). After washing and pre-incubating overnight in α-MEM medium supplemented with 2 μM cilostamide, oocytes were released in inhibitor-free medium for *in vitro* maturation at 37 °C under 5% CO_2_.

### Oocyte morpholino antisense oligonucleotide microinjections

Germinal vesicle (GV) stage oocytes were isolated from wild type (WT) or Dazl Heterozygous (Dazl^+/−^) mice. After pre-incubated in α-MEM medium supplemented with 2 μM cilostamide for 1hr at 37°C under 5% CO_2_, 5–10 pl of 1 mM morpholino oligonucleotides (Gene Tools) of standard control (5’-CCT CTT ACCT CAGTT ACAATTTAT A-3’) or against Dazl (5’- CCTCAGAAGTTGTGGCAGACATGAT-3’) were injected into WT or Dazl^+/−^ oocytes using a FemtoJet express microinjector. Following overnight incubation in α-MEM containing 2 μM cilostamide medium, oocytes were released in α-MEM medium without inhibitor for *in vitro* maturation or recording under the microscope.

### RiboTag IP RNASeq

Oocytes from RiboTag wild type and *Dazl*^+/−^ mice were collected as described above. Wild type oocytes were injected with a CON-MO, while the *Dazl*^+/−^ oocytes were injected with a DAZL-MO. Control experiments show that *Dazl*^+/−^ oocytes have maturation timing and PB extrusion rates identical to wild type oocytes but a 50% decrease in *Dazl* mRNA and protein. In addition, pilot experiments showed a dosage effect in DAZL depletion and MII stage block when comparing DAZL MO injected in wild type oocytes versus DAZL-MO injected in *Dazl*^+/−^ oocyte.

Oocytes injected with Con-MO and DAZL-MO were precinubated overnight in the presence of 2 uM milrinone and the following morning transferred to maturation medium and incubated for 6 hrs. At the end of the incubation, only oocytes that had undergone GVBD were collected in 5 μl 0.1% polyvinylpyrrolidone (PVP) in PBS, flash frozen in liquid nitrogen, and stored at −80°C. In parallel, GV oocytes were kept in milrinone, then harvested and processed together with the MI oocytes. A total of 2000 oocytes (0 hr and 6 hrs with either CON-MO or DAZL-MO injection) were injected and cultured for the duplicate determination of the effect of DAZL depletion on ribosome loading of endogenous mRNAs. On the day when the RiboTag IP was performed, oocytes were thawed, lysed and an aliquot of the oocyte extract was saved and stored to measure total transcript levels before the IP. RiboTag IP was performed as described in the section on Immunoprecipitation. After IP, all samples were used for RNA extraction using the RNeasy Plus Micro kit (Qiagen, 74034). The quality of the extracted RNA was monitored with Bioanalyzer chips (Agilent). RNA samples were transferred to the Gladstone Institutes Genomics Core for cDNA library preparation using the Ovarion RNA-Seq System V2 (NuGen). Samples were sequenced using the HiSeq400 platform.

### Real-time qPCR

Real-time qPCR was performed using Power SYBR PCR master mix with ABI 7900 Real-Time PCR system (Applied Biosystems). All oligonucleotide primers used in this project were designed against two exons flanking an intron to avoid amplification of genomic DNA (Supplementary Table 1). The specificity of each pair of primers was verified by using a unique dissociation curve, performed at the end of the amplification. Data was normalized to its corresponding input and IgG in RiboTag/qPCR for HA and DAZL antibody and expressed as the fold-enrichment of 2^-ΔΔCt^.

### Western Blotting

Oocytes were lysed in 10μl 2x Lammli buffer (Bio-Rad) supplemented with mercaptoethanol, and a cocktail of phosphatase and protease inhibitors (Roche). The oocyte lysates were boiled for 5 mins at 95 °C and then transferred to an ice slurry, then separated on 10% polyacrylamide gels and transferred to a polyvinylidene difluoride (PVDF) membrane. Membranes were blocked in 5% milk for 1 hr at room temperature and incubated with primary DAZL antibody (ab215718, Abcam, 1:1000) overnight at 4 °C. An antibody against α-tubulin (T6074, Sigma-Aldrich; 1: 1000) was used as a loading control. After overnight incubation, membrane was washed in TBS-Tween 20 (0.05%) three times and incubated with HRP-conjugated secondary antibodies (Pierce; 1:5000) for 1 h at room temperature. The signals were detected using Super Signal West Fremto (Thermo Scientific, 34095).

### Culture of ES Cells

ES cells were handled under sterile conditions and recovered in DMEM medium (high glucose, Glutamax, Pyruvate) at room temperature. After centrifuging the cells at 200x g and resuspending with 1.5 ml DMEM culture medium supplemented with 15% KOSR, 2% FBS, 1% 2-Mercaptoethanol, 1% penicillin/streptomycin, 1% MEM Non-Essential Amino Acids, 1000 U/ml LIF, 3 μM CHIR-99021, and 1 μM PD0325901, ES cells were cultured in multi-well plates coated with 0.1% gelatin in H_2_O and FBS at 5% CO**2** and 37°C. Culture media was changed daily. When ES cells reached approximately 80% confluency, they were washed once with DPBS and incubating with 0.05% Trypsin for 1 min. The reaction was stopped by adding ES cell culture medium without LIF and 2i and KOSR, KOSR has been replaced with 10% FBS in this medium (later called MEF Media). Cells were then pelleted by centrifuging at 200 x g for 5 min. The ES cell pellet was dissolved in RNase-free PBS and stored at −80°C for DAZL IP.

### Immunoprecipitation

RiboTag IP or DAZL RIP analysis was performed as described previously^18, 19^. Briefly, GV-arrested or MI oocytes (200 oocytes/sample) were washed and collected in RNase-free PBS with 1% polyvinylpyrrolidone. After lysis in 300 μl of supplemented homogenization buffer S-HB; 50 mM Tris-HCl pH 7.4, 100 mM KCl, 12 mM MgCl2, 1% NP-40, 1 mM dithiothreitol, protease inhibitors, 40 U RNAseOUT, 100 μg/ml cycloheximide and 1 mg/ml heparin (Sigma-Aldrich, H3393)]., samples were centrifuged at 12,000 g for 10 mins and supernatants were precleared with prewashed Protein G magnetic Dynabeads (Invitrogen, 10007D) for 30 min at 4°C. 15μl of precleared lysates was aliquot for input (total transcripts) and stored at −80°C for mRNA extraction in next day. The remain precleared lysates were incubated with specific antibody (anti-HA antibody, anti-DAZL antibody) or its corresponding IgG (mouse IgG, ab37355; rabbit IgG, ab37415; Abcam) 4 hrs at 4°C on a rotor. Then pre-washed Protein G magnetic Dynabeads were added in the lysates for overnight incubation at 4°C on a rotator. The following day, bead pellets were washed three times in 500 μl homogenization buffer (HB) on a rotor at 4°C for 10 min. Two more washes were performed with 1M urea/high-salt buffer for 10 min each. RNA eluted from beads was either HA-tagged ribosome associated transcripts or IgG (no-specific binding transcripts), together with input for extraction. In some experiments, the specificity of the immunoprecipitation was determined by using WT rather than RiboTag mice. RNA was purified with RNeasy Plus Micro kit (Qiagen, 74034) according to manufacturer’s instructions and directly used for RNA-Seq analysis or reverse transcription for qPCR analysis. cDNA was prepared using SuperScript III First-Strand Synthesis system (Invitrogen, 18080-051) with random hexamer oligonucleotide primers. cDNA samples were stored −80°C for following experiments.

For the RiboTag/qPCR analysis, *Ccnb1, Dppa3* and *Gdf9* (transcripts not regulated by DAZL as previous reported^18, 19^), were used to normalize the qPCR data. Zp3 contains no recognizable DAZL-binding element and was used as negative controls for DAZL immunoprecipitation. The data are reported as fold enrichment, with IgG values set to 1.

### Reporter mRNA preparation and reporter assay

The *Oosp1, Obox5, CenpE* and *Ccnb1* 3’UTR sequences were obtained by sequencing oocyte cDNA and cloned downstream of the YPet coding sequence. An oligo (A) stretch of 20A was added in each construct. All constructs were prepared in the pcDNA 3.1 vector containing a T7 promoter, allowing for in vitro transcription to synthesize mRNAs, and fidelity was confirmed by DNA sequencing. mRNA reporters were transcribed *in vitro* to synthesize mRNAs with mMESSAGE mMACHINE T7 Transcription Kit (Ambion, AM1344); polyadenylation of *mCherry* was obtained using Poly(A) Tailing Kit (Ambion, AM1350). All the messages were purified using MEGAclear Kit (Ambion, AM1908). mRNA concentrations were measured by NanoDrop and its integrity was evaluated by electrophoresis.

Time-lapse recordings were performed using a Nikon Eclipse T2000-E equipped with mobile stage and environmental chamber of 37°C and 5% CO_2_. *YFP-Oosp1, YFP-Obox5, YFP-Cenpe* or *YFP-CcnB1* were injected at 12.5 ng/μL with either CON-MO or DAZL-MO. Each *YFP-3’UTR* reporters we also co-injected with polyadenylated *mCherry* at 12.5 μg/μL in oocyte. After injection, oocytes were pre-incubated overnight in α-MEM medium supplemented with 2 μM cilostamide to allow expression of the reporters. mCherry signals did not change significantly in oocytes at different stages of maturation. Ratios of YFP reporter and the level of mCherry signal measured at plateau in each oocyte were calculated. In those cases where DAZL ablation had an effect in GV oocytes, the data were normalized to the signal of GV stage accumulation of corresponding proteins. Rate of translation associate with reentering into cell cycle (after GVBD versus before GVBD) were calculated by fitting YFP:mCherry data and calculating the slope of the interpolation obtained by linear regression (Prism) prior to GVBD or after GVBD when a new rate of translation had stabilized.

### DAZL RIP-Chip analysis

DAZL RIP-Chip was performed as previously reported ^18^. Briefly, C57BL/6 female mice (22–24 days old) were primed with PMSG, after 48 hrs, mice were stimulated with hCG for 0 hr, 6 hrs, or 14 hrs to collect GV, MI, and MII stage oocytes. Oocyte lysates were centrifuged at 12,000 g for 10 mins at 4°C. Supernatants were used for RNA extraction. RNA was purified with RNeasy Plus Micro kit (Qiagen). RNAs in the RNP fractions were reverse-transcribed with SuperScript III (Invitrogen). Five micrograms of cDNA was fragmented and hybridized with Affymetrix Mouse Genome 430.2 array chips^46^. DNA-Chip Analyzer (dChip) was used for normalization and to quantify microarray signals with default analysis parameters. Comparison between samples was performed using dChip with a fold change of 1.5, FDR < 5%, and P < 0.05.

### Statistical analysis

Statistical analysis was performed using the GraphPad Prism8 package. The statistical analysis performed depended on specific experiment and is reported in the figure legend. For comparison between two groups, two-tailed paired t-test was used. Statistical significances is denoted by asterisk in each graph. The quality check of RNA-Seq reads was performed using FastQC and reads were then trimmed with Trimmomatic. Alignment of the reads to the mouse genome was performed by Hisat2, .bam files were sorted and indexed using Samtools, and count files were generated by HTSeq. TMM normalization and the remaining RNA-Seq statistical analyses were done through edgeR and other Bioconductor scripts.

## Acknowledgements

We wish to thank Dr. Matthew Cook for the help in the RIP-Chip experiments. We thank Dr. Soeren Muller and Dr. Xiaoyuan Zhou for their help and advice on processing and analyzing the RNA-Seq data. This work was supported by NIH R01 GM097165, GM116926 and the Specialized Cooperative Centres Programme in Reproduction and Infertility Research (P50HD055764-10), Eunice Kennedy Shriver National Institute of Child Health and Human Development (NICHD) to MC. Enrico Maria Daldello is supported by a Fellowship from the Lalor Foundation. Xuan G. Luong is supported by a T32-HD007263 Integrated Training in Reproductive Sciences.

