## Supplemental information for "The RNA binding protein DAZL functions as repressor and activator of maternal mRNA translation during oocyte maturation"

*Cai-Rong Yang et al.*

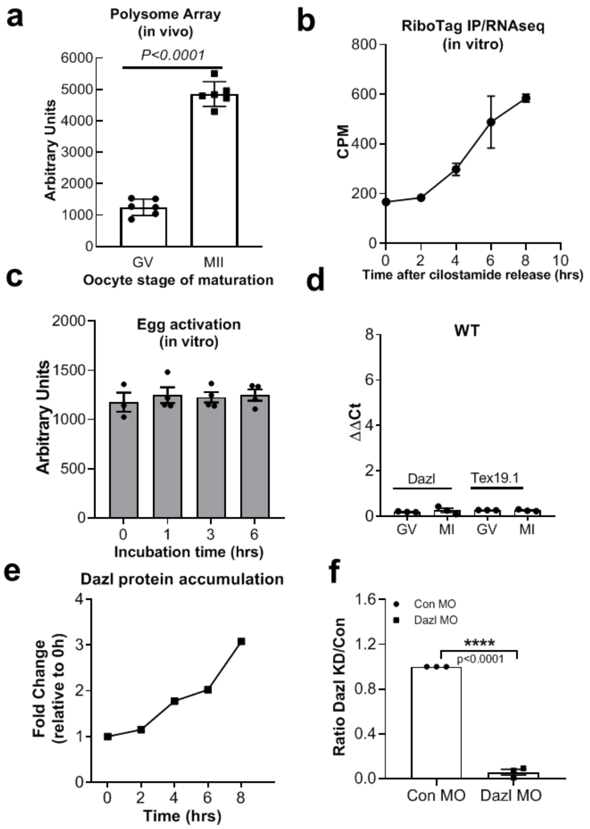


**Supplementary Figure 1 Dazl mRNA translation is regulated during oocyte maturation.**

**(a)** The Dazl mRNA becomes associated with the polysome fraction at the GV-to-MII transition *in vivo*. Polysome array data described in previous publications were mined for the Dazl mRNA association with polysomes. The data are the Mean + SEM of six independent biological replicates. **(b)** Ribosome loading of *Dazl* mRNA is increased during in vitro oocyte maturation. After harvesting, oocytes were incubated *in vitro* for the times reported in the abscissa. At the end of the incubation, samples were processed for RiboTag IP/RNAseq as described in the 'Materials and Methods'. The data are the mean +/- range of duplicate samples for each time point. The details of the experiments are summarized in a paper in preparation. **(c)** Polysome associated transcript of *Dazl* is not changed upon egg activation in vitro. MII oocytes were treated *in vitro* Sr2+ for the indicated times. At the end of the incubation, cells were harvested and processed for polysome arrays as described. Each point is the Mean +/- SEM of three or four biological replicates. **(d)** RiboTag IP/qPCR of oocyte extracts from wild type mice. The mRNA recovered is at background levels and is similar in all the experimental groups. **(e)**Protein levels measurement for DAZL accumulation during oocyte maturation until 8 hrs (MI stage). **(f)** Measurement of protein levels for DAZL expression with DAZL-KO MO injection. Each group is the average and SEM of three biological replicates.

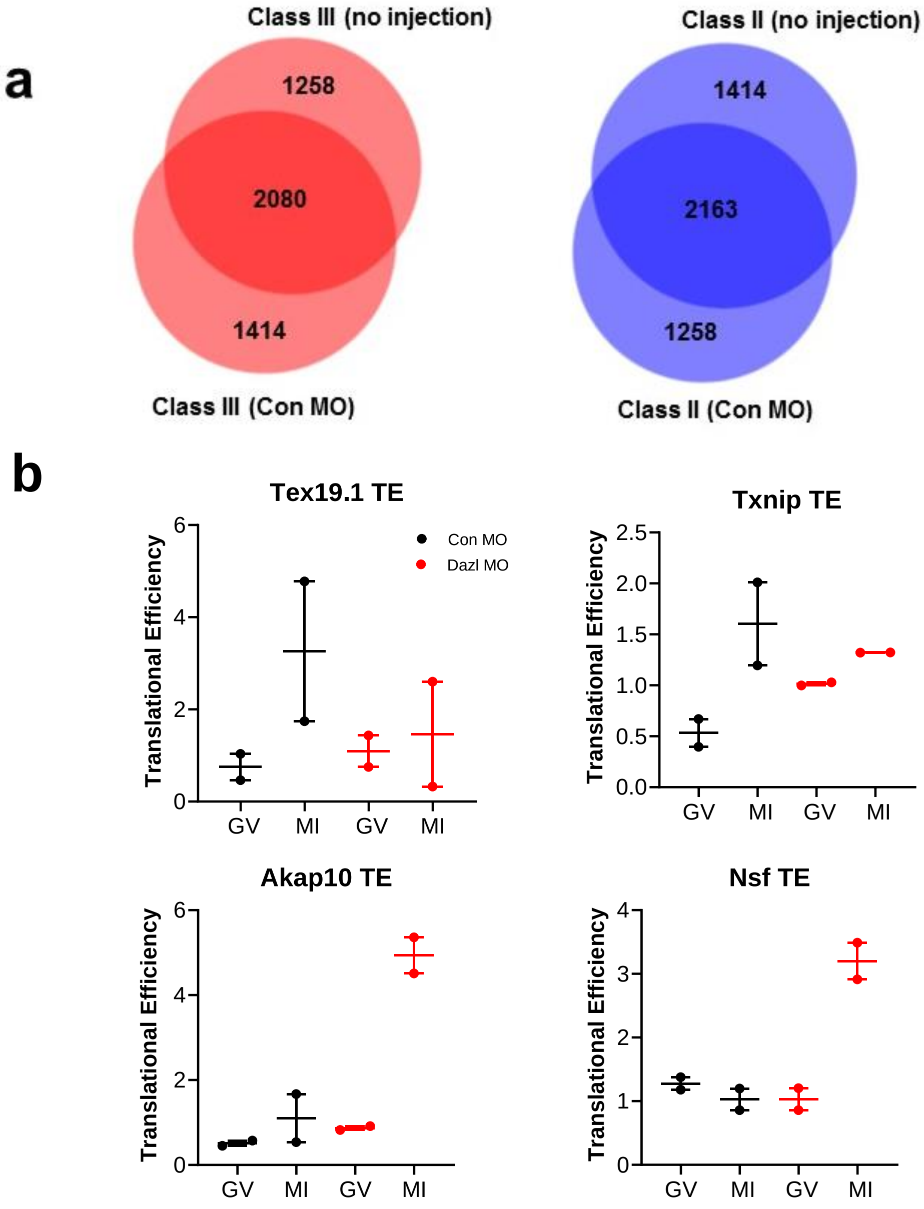


### Supplementary Figure 2

**(a)** Comparison of the RiboTag IP/RNAseq data from CON-MO injected oocytes with RiboTag IP/RNASeq data for non-injected oocytes. Increased and decreased ribosome loading onto mRNAs at 6 hrs is compared. Qualitatively, the regulation of translation is comparable between the two experiments for the majority of transcripts. Quantitatively the data could not be directly compared because of batch effects. **(b)** Calculation of the translational efficiency (TE) for the candidate mRNAs reported in Fig. 2C. TE is calculated as the CPM of ribosome bound mRNA divided by the total mRNA levels expressed in CPM.

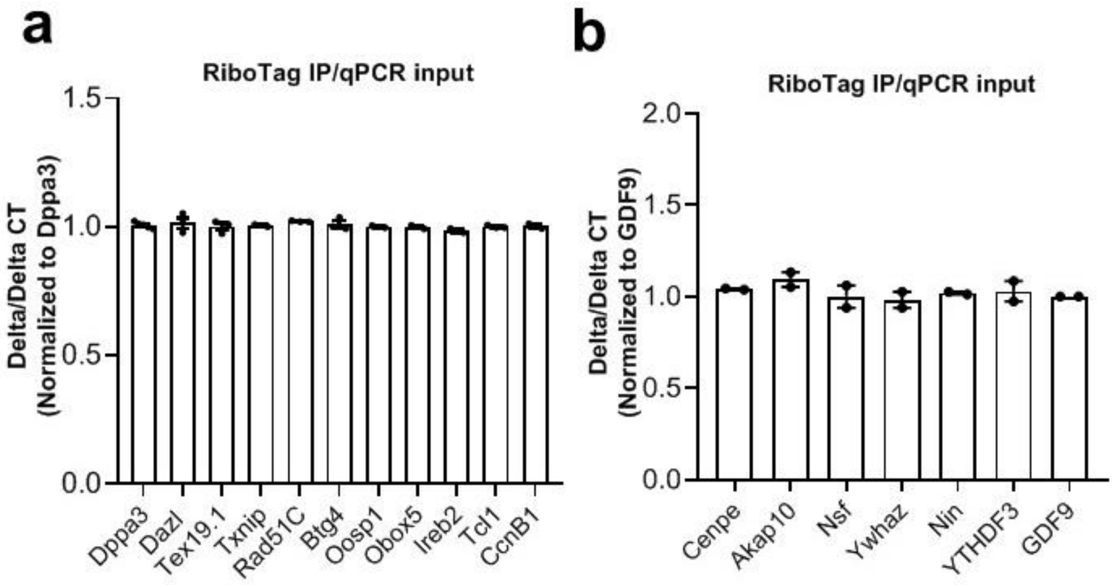


**Supplementary Figure 3**

Total mRNA levels of the representative Dazl targets reported in Figure 3. GV stage oocytes from wild type or Dazl$^{+/-}$ mice were injected with CON-MO or DAZL-MO. After overnight preincubation with 2 µM milrinone, oocytes were cultured in inhibitor-free medium for maturation. Oocytes were collected at 6 hrs for qPCR analysis of total transcript levels. The samples are the same as those reported in Figure 3.

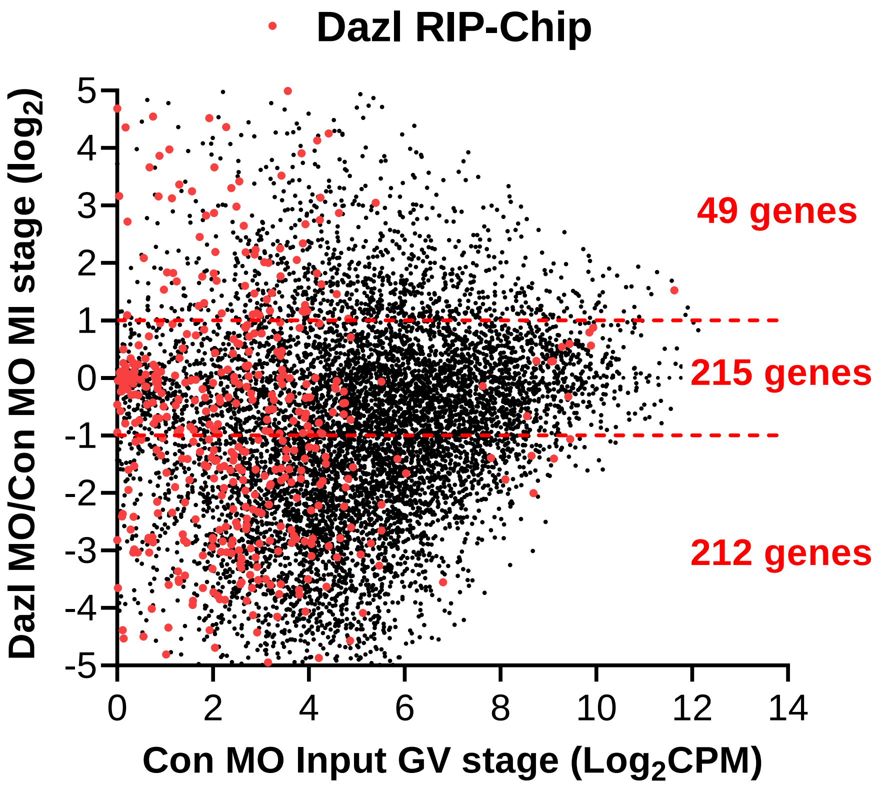


**Supplementary Figure 4**

Comparison of genes whose ribosome loading is affected by DAZL depletion and their interaction with DAZL protein. The plot reports the ratio of log2 fold changes of RiboTag IP/RNASeq between the DAZL-MO and CON-MO. The genes marked in red are specifically immunoprecipitated ($P<0.05$; >1.5 over IgG) by a DAZL antibody in the DAZL RIP-Chip experiment.  The dashed red lines mark the 2fold change threshold.

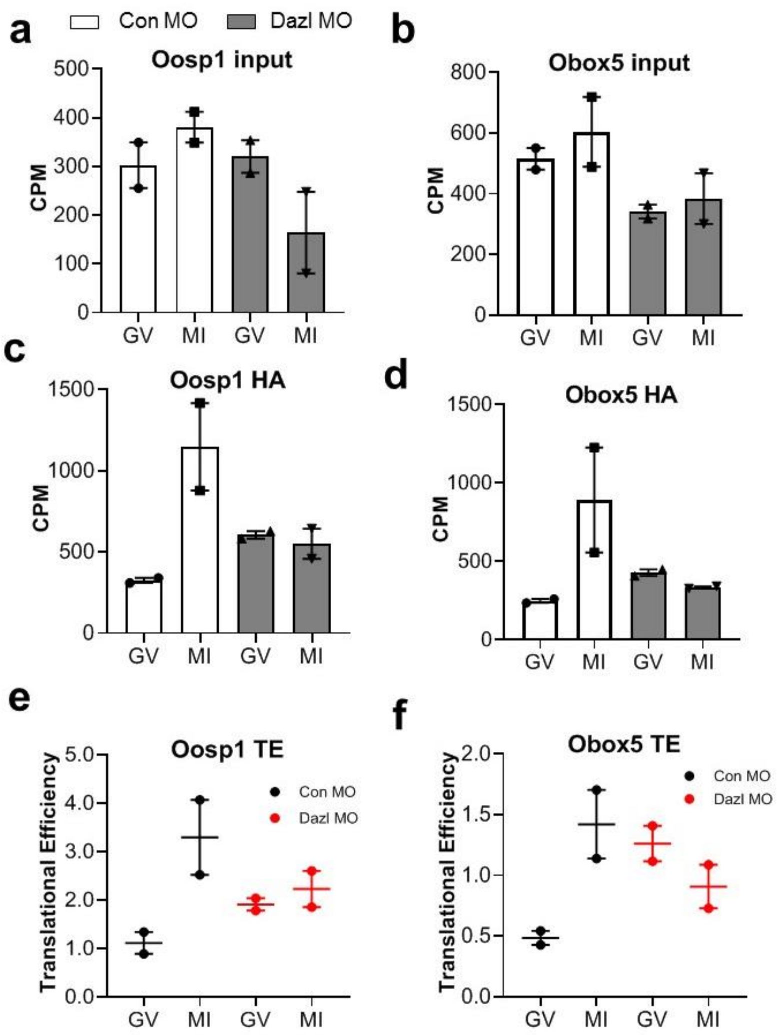


**Supplementary Figure 5**

RiboTag IP/RNASeq data and TE calculation for the *Oosp1* and *Obox5* mRNAs. The data are calculated as described above. Notice the loss of translational activation in oocytes depleted of DAZL. This is due to increased levels of translation during GV-arrested and decreased activation during MI.

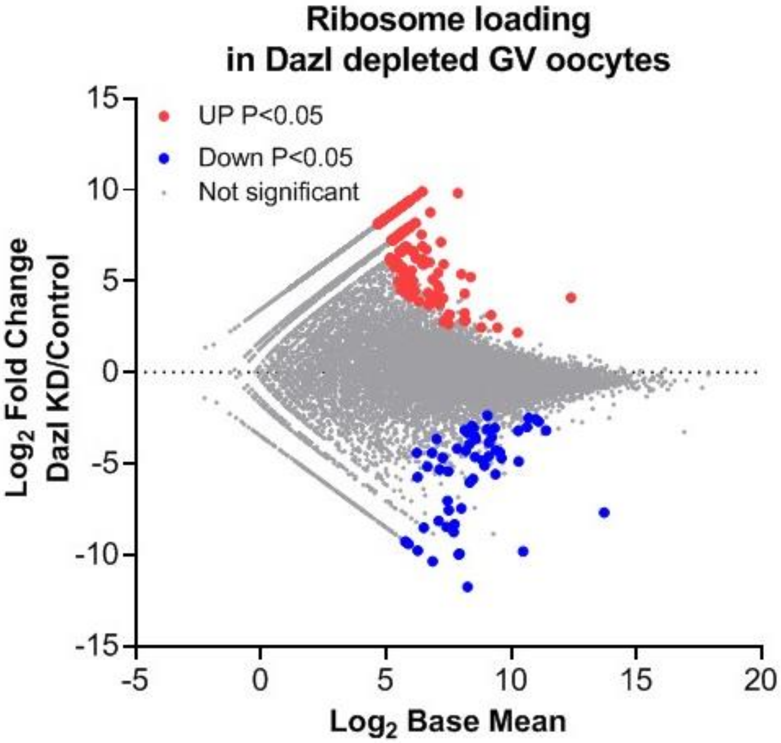


**Supplementary Figure 6.**

Effect of DAZL depletion on ribosome loading of mRNAs in GV-arrested oocytes. The ribosome loading of 153 transcripts was significantly increased in GV oocytes depleted of DAZL whereas 69 transcripts were decreased (no fold change cutoff, FDR < 0.05).

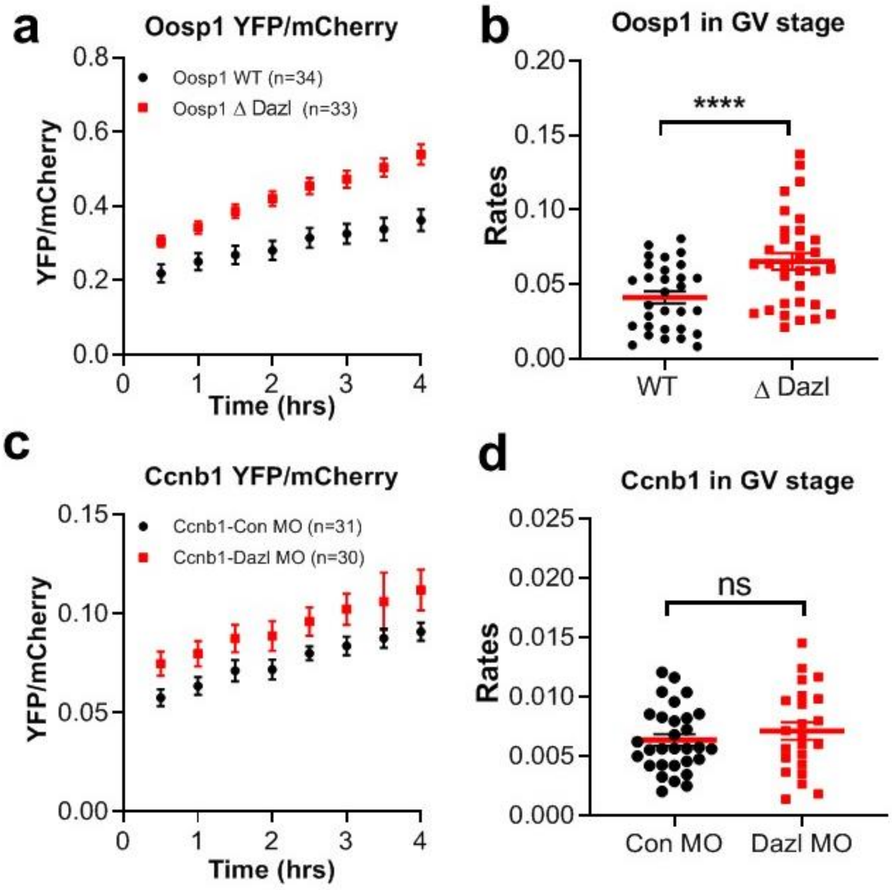


**Supplementary Figure 7. Oosp1 accumulation during overnight incubation**

Mutation of the DAZL-binding element causes an increase in translation rate of the *YFP-Oosp1* reporter in GV-arrested oocytes. Oocytes were injected with the YFP reporter fused wild type or mutated DAZL-binding element 3'UTR of *Oosp1*. After 3 hrs of preincubation for recovery, the oocytes were maintained in GV with cilostamide and florescence monitored by time lapse microscopy. The number of oocytes analyzed in each group is reported in the figure.

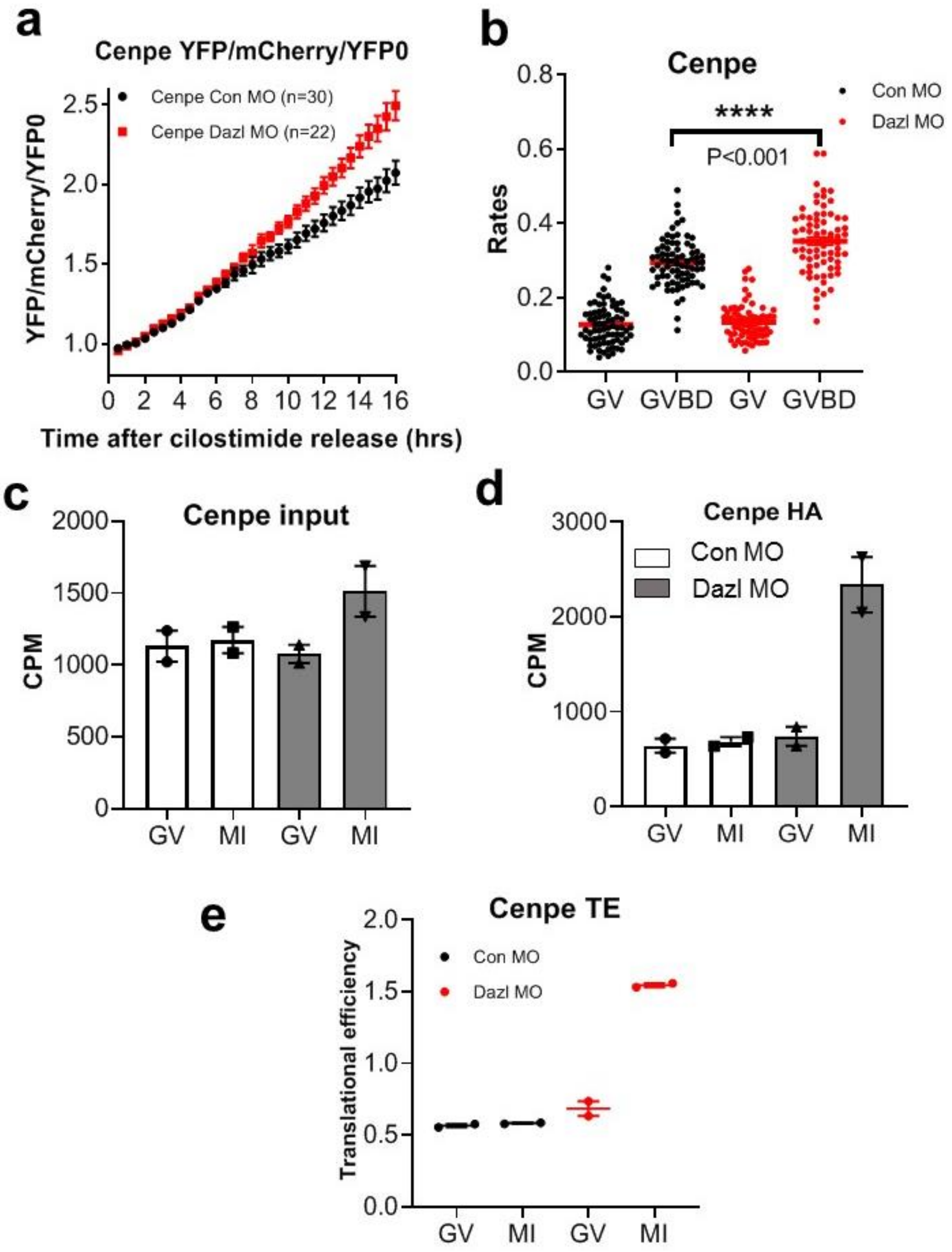


**Supplementary Figure 8**

Injection of the *YFP-CenpE* reporter construct recapitulates the effect of DAZL depletion on ribosome loading of the endogenous *CenpE* mRNA. **(a and b)** Accumulation of the YFP-CenpE reporter during oocyte maturation and measurement of rates of translation in individual injected oocytes. **(c and d)** RiboTag IP/RNASeq data for the endogenous *CenpE* mRNAs. Each point is the average and range of two biological replicates.

### Supplementary Table 1

| Name | Primers |
|---|---|
| Akap10 FW | AGTCATCAGATTCCCACCGAC |
| Akap10 REV | TGGCTTCTGTAATTGGTATTGGC |
| Cenpe FW | TGTCTGTGTTCGTGTGCGAC |
| Cenpe REV | AAGATTTACCCCCATCGCTCT |
| Ywhaz FW | AAAGGCAGGGCGTCATTCAG |
| Ywhaz REV | ACGGGGTTTCCTCCAATCAC |
| Nin FW | AGAACTCTATCCAGTGAGGAGC |
| Nin REV | TAGTGGCTCAAGCACTGTCAC |
| Dazl FW | GGATGAAACCGAAATCAGGA |
| Dazl REV | ATAGCCCTTCGACACACCAG |
| Oosp1 FW | TCTCTGGGGTTTGATCTTCAGC |
| Oosp1REV | CGTAGGCCTGATCCTTAGATGG |
| Obox5 FW | AGGGGATATCATGTTGGAGCC |
| Obox5 REV | GTTCCTTGCCGGTTCTTGAG |
| Tex19.1 FW | GGCTGTACCATCTTGTCCA |
| Tex19.1 REV | TCCTCTTCCTCTTCCTCCTC |
| Txnip FW | TGGACGACTCTCAAGACAGC |
| Txnip REV | CATTTCCTGCAGGCTCACTG |
| Tcl1 FW | TCGGAGTCCAACGATGAATAACC |
| Tcl1 REV | CTTCTTGGAGCCCAGTGTAGAG |
| Btg4 FW | TCGATCCCTATGAGGTGTGC |
| Btg4 REV | CCTGCTGCAGCTTTCTTCATC |
| Rad51C FW | TCAAGCTTTGCTTGTTCCATTA |
| Rad 51C REV | TATCGTAGACTCCTTCTGGCTTG |
| Ireb2 FW | TCAATGTACCTAAACTTGGCGG |
| Ireb2 REV | AAGGGCACTTCAACATTGCTC |
| Gdf9 FW | CAAACCCAGCAGAAGTCAC |
| Gdf9 REV | TTAAACAGCAGGTCCACCA |
| Ccnb1 FW | AAGGTGCCTGTGTGTGAACC |
| Ccnb1 REV | GTCAGCCCCATCATCTGCG |
| Dppa3 FW | GACCCAATGAAGGACCCTGAA |
| Dppa3 REV | GCTTGACACCGGGGTTTAG |
| Zp3 FW | TTTCGGCATTTCAAGTCCC |
| Zp3 REV | GGTGATGTAGAGCGTATTTCTG |
| Oosp1 3’UTR FW | CATCACCATTGAatggtctggtgatttcttatctcc |
| Oosp1 3’UTR REV | GCGGGTTTAAACttagacacggcactaatggg |
| Oosp1 YFP FW | gtgccgtgtctaaGTTTAAACCCGCTGATCAGCCTC |
| Oosp1 YFP REV | tcaccagaccatTCAATGGTGATGGTGATGATGAC |
| Obox5 3’UTR FW | CATCACCATTGAcatatcagatgactggcttac |
| Obox5 3’UTR REV | GCGGGTTTAAACaaagaaatttaaatttactattttctcc |
| Obox5 YFP FW | ttaaatttctttGTTTAAACCCGCTGATC |

| | |
|---|---|
| Obox5 YFP REV | tcatctgatatgTCAATGGTGATGGTGATG |
| Cenpe 3'UTR FW | CATCACCATTGAatgcccctgtcccgtc |
| Cenpe 3'UTR REV | AGCGGGTTTAAACccttcaagacctttattcttcgc |
| Cenpe YFP FW | aaaggtcttgaaggGTTTAAACCCGCTGATCAGCCTC |
| Cenpe YFP REV | ggacaggggcatTCAATGGTGATGGTGATGATGAC |
